# An androgen-derived transcriptional state reveals signaling and mitochondrial vulnerabilities in cancer

**DOI:** 10.64898/2026.10.06.757186

**Authors:** Yingqi Miao, Takashi Baba

## Abstract

Transcriptional states can provide functional information beyond genomic alterations, but whether states derived from nonmalignant physiological perturbations remain informative in cancer is unclear. We separated global RNA-output suppression from relative transcriptional remodeling induced by dihydrotestosterone in mouse adrenal zona fasciculata cells and derived a cross-sex transcriptional signature. Projection of this signature to human orthologs defined an androgen-suppressed transcriptional state (ASTS) across cancer cell lines. Integration with pharmacogenomic and CRISPR dependency datasets showed that higher ASTS was associated with resistance to EGFR-targeted drugs and reduced EGFR dependency, together with increased dependency on mitochondrial pathways, including mitochondrial translation and respiratory electron transport. These associations were not attributable to a single major cancer lineage and remained evident after adjustment for canonical oncogenic drivers and cell-state covariates. ASTS provides a proof of concept that an endocrine-derived transcriptional state can reveal candidate functional vulnerabilities in an unrelated cancer context.

## Introduction

Precision oncology has been shaped by the identification of genomic alterations that define oncogenic dependencies and therapeutic opportunities. However, large-scale pharmacogenomic studies have shown that responses to anticancer agents are associated not only with specific genomic alterations but also with cancer lineage and gene-expression features^1–4^. Functional-genomic studies have similarly demonstrated that transcriptional information can be highly informative for predicting genetic dependencies; in an early Cancer Dependency Map, expression-based features represented the strongest predictors for the majority of dependencies for which predictive models could be constructed^5^. Large-scale CRISPR studies have further mapped selective cancer dependencies and prioritized therapeutic targets^6,7^. More recently, transcriptionally defined cancer cell states have been used directly to identify selective vulnerabilities. For example, a mesenchymal-like state was used to identify PKN2 as a dependency across diverse cancer lineages^8^. Together, these observations support the concept that transcriptional state provides a layer of functional information that can complement genomic characterization in the search for therapeutic vulnerabilities. A further example is the dependence of particular mesenchymal and therapy-resistant states on GPX4^9^.

Gene-expression signatures offer a means of representing and transferring such states across experimental systems. The Connectivity Map established that transcriptional responses to defined perturbations can uncover functional relationships among drugs, genes, and biological states, and the L1000 framework subsequently extended this approach to large-scale perturbational profiling^10,11^. Most applications relevant to cancer have focused on signatures derived from malignant cells, treatment responses, or disease-associated transcriptional programs. It remains less clear whether a transcriptional state derived from a defined endocrine perturbation in nonmalignant tissue can retain functional information when transferred across species and into an unrelated cancer-cell context. Such transfer would provide a way to exploit experimentally defined biological states as probes of functional-genomic datasets rather than requiring the state itself to originate from cancer.

The adrenal zona fasciculata (zF) provides a useful system in which to address this question because androgen alters both transcriptional output and cellular metabolism in a defined in vivo context. We previously showed that androgen suppresses global gene expression in zF cells, in association with reduced expression of Ad4BP/SF-1 (NR5A1)^12^, a transcription factor that broadly regulates adrenal gene expression and metabolic function. Earlier studies also demonstrated that Ad4BP/SF-1 coordinates metabolic programs including glycolysis and cholesterol biosynthesis^13,14^. External spike-in normalization distinguishes changes in global RNA abundance from relative transcriptional remodeling^15^. In the zF system, this distinction applies to the previously reported DHT-associated reduction in RNA output. This distinction suggested that the androgen response could be considered as two related but conceptually separable features: a change in total transcriptional output and a pattern of relative transcriptional remodeling among genes. We asked whether the latter contains biological information that remains informative beyond the adrenal context in which it was defined.

Here, we reanalyzed cell-number-normalized mouse zF RNA-sequencing data to distinguish DHT-associated global RNA-output suppression from relative transcriptional remodeling. We derived a cross-sex signature of the relative DHT response, projected the signature to one-to-one human orthologs, and used within-sample expression ranks to define an androgen-suppressed transcriptional state (ASTS) score across human cancer cell lines. We then integrated ASTS with separately generated pharmacogenomic and genome-scale CRISPR dependency datasets to ask whether this mouse-derived state was associated with functional phenotypes in human cancer cells. ASTS was associated with resistance to EGFR-targeted drugs and reduced EGFR dependency, together with a selective dependency on mitochondrial functions. These findings provide a proof of concept that a transcriptional state derived from a defined in vivo endocrine perturbation can retain functional information after transfer to a distinct cancer-cell context and suggest that transcriptional state information can complement genomic characterization when prioritizing candidate therapeutic vulnerabilities.

## Methods

### Study design

This study reanalyzed previously generated cell-number-normalized RNA sequencing (CNN RNA-seq) data from mouse adrenal zona fasciculata (zF) cells and integrated the resulting transcriptional signature with publicly available transcriptomic, pharmacogenomic, and CRISPR dependency datasets from human cancer cell lines. The analysis comprised four major steps: separation of global and relative transcriptional responses to dihydrotestosterone (DHT) in mouse zF cells; construction and mouse-to-human projection of a DHT-associated relative transcriptional signature; quantification of the resulting androgen-suppressed transcriptional state (ASTS) in human cancer cell lines; and association of ASTS with drug response and genetic dependency. The mouse-derived signature was fixed before the pharmacogenomic analyses were performed.

### Mouse adrenal zF CNN RNA-seq dataset

The mouse adrenal zF CNN RNA-seq dataset was generated previously using Ad4BP-BAC-EGFP transgenic mice. zF cells were purified on the basis of EGFP expression, and RNA isolated from 10,000 cells per sample was used for RNA sequencing without normalization to total cellular RNA. Thus, the dataset retains information on differences in RNA abundance per fixed cell number. The experimental groups consisted of sham-operated males, orchiectomized males treated with oil, DHT, or estradiol (E2), sham-operated females, and ovariectomized females treated with oil, DHT, or E2, with three biological replicates per group. The generation of the mice, gonadectomy, hormone administration, zF purification, and original CNN RNA-seq procedures were described previously^12^.

The original CNN RNA-seq datasets were deposited in the DNA Data Bank of Japan under accession numbers DRA015952 and DRA017476.

For the present study, the primary DHT contrasts compared DHT-treated with oil-treated orchiectomized males and DHT-treated with oil-treated ovariectomized females. Sham-operated and E2-treated groups were used for additional biological and specificity analyses.

No new animal experiments were performed in the present study; all animal experiments underlying the previously generated RNA-seq datasets were approved by the Animal Care and Use Committee of Kyushu University and were performed in accordance with institutional guidelines, as described previously^12^.

### Read counting and RNA-seq normalization

Trimmed paired-end reads were aligned to a custom mouse reference based on mm10 and supplemented with External RNA Controls Consortium (ERCC), EGFP, and mCherry sequences using STAR version 2.7.11b^16^. Gene-level counts were generated from coordinate-sorted BAM files using featureCounts in Subread version 2.1.1^17^. The primary count matrix was generated using exon features grouped by gene ID, with multimapping and overlapping reads retained (-M -O), paired-end fragments counted (-p --countReadPairs), and reverse-strand specificity applied (-s 2). An otherwise identical unstranded count matrix (-s 0) was generated as an alternative counting workflow for comparison. The strand-aware matrix was used for downstream analyses.

Two normalization strategies were applied using DESeq2 version 1.46.0 in R version 4.4.3^18^. Standard DESeq2 size factors were estimated from the endogenous count matrix across all 24 samples using the median-of-ratios method. In parallel, ERCC-based normalization was used to preserve differences in endogenous RNA abundance relative to the exogenous spike-in controls. For each sample, the ERCC size factor was calculated as the total ERCC count divided by the geometric mean of ERCC totals across all 24 samples. Both differential expression analyses used raw endogenous counts as input to DESeq2, with either the standard or ERCC-derived size factors assigned to the model. ERCC-normalized counts were obtained by dividing endogenous counts by the corresponding ERCC size factor. ERCC-based normalization was used to assess the global RNA output component of the response, whereas standard DESeq2 normalization was used to quantify relative gene-specific remodeling.

For the normalization comparison, genes with a total count below 10 across all 24 samples were excluded, retaining 16,535 genes. For each normalization strategy, a DESeq2 model with experimental group as the explanatory variable was fitted to all 24 samples, and male and female DHT contrasts were extracted from that model. For ASTS construction, the dataset was restricted to the 12 male and female gonadectomized oil- and DHT-treated samples. Applying the same total-count threshold of 10 across these 12 samples retained 15,729 genes. The standard DESeq2 size factors estimated from the full 24-sample dataset were retained for the 12-sample ASTS models.

The strand-aware and alternative unstranded counting workflows were compared using the same samples. For each sample, agreement between workflows was evaluated across genes using Pearson correlation of log1p-transformed counts and Spearman rank correlation of counts (Fig. S1A).

### Definition of DHT-responsive genes and construction of the mouse ASTS signature

Relative DHT effects were estimated using a pooled additive DESeq2 model with sex and DHT treatment as explanatory variables and the standard DESeq2 size factors described above. Genes were classified as broadly DHT responsive when the pooled DHT effect had an adjusted P value < 0.05 and the male and female DHT effects were each in the same direction as the pooled effect. The cross-sex core additionally required an absolute pooled log2 fold change >= 0.25 and an absolute sex-specific log2 fold change >= 0.1 in each sex. Genes with a strong sex-by-DHT interaction, defined by an adjusted P value < 0.05 and an absolute interaction log2 fold change >= 0.5, were excluded from the core. Interaction effects were estimated using a DESeq2 model including sex, DHT treatment, and their interaction. These criteria yielded 4,356 broadly DHT-responsive genes and 3,259 cross-sex core genes, comprising 1,723 DHT-relative-up and 1,536 DHT-relative-down genes.

Within the cross-sex core, DHT-relative-up and DHT-relative-down genes were ranked separately in descending order of the absolute Wald statistic for the pooled DHT effect. Ties were resolved by descending absolute pooled log2 fold change, ascending adjusted P value, and gene identifier. The top 50 genes in each direction were selected to generate the frozen 100-gene mouse ASTS signature.

E2 responses were examined subsequently as a specificity analysis and were not used to select the signature. Cross-sex core genes were classified relative to the pooled DHT-effect direction. Genes were classified as E2-concordant when the E2 effects in both sexes were in the DHT-defined direction and the E2 response had an adjusted P value < 0.05 in at least one sex. Genes were classified as E2-opposite when neither sex-specific E2 effect was in the DHT-defined direction and the E2 response had an adjusted P value < 0.05 in at least one sex. Genes with no significant E2 response in either sex were classified as nonsignificant; the remaining genes were classified as mixed/weak (Fig. S2A,B).

### Mouse-to-human orthology mapping

The frozen 100-gene mouse signature was mapped to human genes using Ensembl release 114 mouse-to-human orthology annotations^19^. One-to-one orthology was the prespecified criterion for inclusion in the primary human signature. Seventy-eight mouse genes met this criterion, comprising 40 DHT-relative-up and 38 DHT-relative-down genes. Genes with one-to-many or many-to-many mappings, genes without an ortholog in this Ensembl release, and genes whose mouse identifiers could not be resolved unambiguously were excluded; excluded genes were not replaced or re-ranked. One of the 78 one-to-one orthologs carried a lower Ensembl orthology confidence annotation. Because the orthology relationship, rather than the confidence flag, was the prespecified primary inclusion criterion, this gene was retained in the 78-gene primary signature. A 77-gene high-confidence set excluding this ortholog was analyzed separately as a sensitivity set.

### DepMap expression data and calculation of ASTS scores

Human cancer cell-line expression data and model annotations were obtained from DepMap release 26Q1^20,21^, using the OmicsExpressionTPMLogp1HumanProteinCodingGenes.csv expression matrix and the corresponding Model.csv annotations. Expression values were represented as log2(TPM + 1). Only the default expression profile assigned to each ModelID was retained, preventing multiple expression profiles from contributing more than once for the same cell line. After this restriction, 1,719 unique cell lines were included in the primary ASTS analysis. The DepMap portal is available at https://depmap.org/portal; the CCLE molecular-profiling resource provides complementary background on the cell-line expression data^22^.

ASTS was calculated using within-sample ranks to minimize dependence on absolute expression scale. For each cell line, all available protein-coding genes were ranked in ascending order of expression, with average ranks assigned to tied values. Percentile ranks were obtained by dividing each rank by the number of genes with nonmissing expression values. The ASTS score was calculated as the mean percentile rank of DHT-relative-up genes minus the mean percentile rank of DHT-relative-down genes. A positive score therefore indicates that the mean expression rank of the DHT-relative-up genes exceeds that of the DHT-relative-down genes. Higher scores indicate greater resemblance to the relative DHT-associated transcriptional state; ASTS does not estimate absolute RNA output per cell. This use of within-sample ranks and oppositely directed gene sets is conceptually related to single-sample rank-based approaches such as singscore^23^, while the ASTS gene sets and score definition were fixed for the present study.

Scores were calculated independently using the 78-gene primary signature and the 77-gene high-confidence sensitivity set. Agreement between the two scores was evaluated using Spearman rank correlation and the distribution of cell-line-wise score differences.

### Cancer lineage, donor sex, and proliferation-related covariates

Cancer lineage and recorded donor sex were obtained from the corresponding cell-line model annotations.^20^ Cancer lineage was included as a categorical covariate in subsequent transcriptomic, pharmacogenomic, and CRISPR dependency analyses to account for differences among lineages. Lineage annotations followed the OncoTree classification^24^.

A simple proliferation proxy was calculated as the mean within-sample percentile rank of expression for *MKI67*, *PCNA*, *MCM2*, *MCM3*, *MCM4*, *MCM5*, *MCM6*, *MCM7*, *TOP2A*, and *CDK1*. Additional E2F, G2M, combined cell-cycle, and MYC V1 transcriptional scores were constructed independently of ASTS. Genes belonging to either the 78-gene ASTS signature or the proliferation proxy gene set were excluded from these additional scores to minimize direct overlap between predictors.

Associations between ASTS and these scores were evaluated by Spearman rank correlation and by multivariable regression models including cancer lineage, donor sex, the proliferation proxy, and canonical oncogenic driver covariates. For these adjusted score-association models, ASTS, the proliferation proxy, and each outcome score were z-standardized within the analyzed cell lines; HC3 standard errors were used.

### Genome-wide transcriptomic association and gene-set enrichment analysis

For each protein-coding gene in the DepMap expression dataset, log2(TPM + 1) expression was modeled as a function of ASTS, cancer lineage, donor sex, and the proliferation proxy. A driver-adjusted model additionally included the MAPK-, ERBB-, and PI3K-driver variables defined below. Models were fitted by ordinary least squares, and ASTS association t statistics were calculated using conventional model-based standard errors.

Genes were ranked according to the gene-wise ASTS association t statistic. Hallmark and Reactome gene sets from the Molecular Signatures Database (MSigDB; human release 2026.1.Hs)^25^ were obtained using msigdbr version 26.1.1. All 78 genes in the primary ASTS signature were removed from the ranked gene list before enrichment analysis to reduce direct circularity. Gene-set enrichment analysis^26^ was performed using fgseaMultilevel in fgsea version 1.36.2^27^, with gene-set sizes of 15-500 genes, eps = 0, and nPermSimple = 10,000. Transcriptomic and CRISPR GSEA analyses were run in R version 4.5.3 with fgsea version 1.36.2. The primary analysis included 50 Hallmark^28^ and 1,161 Reactome^29^ gene sets. Pathways with non-finite NES, P, or adjusted P values were excluded from the reported enrichment results. Normalized enrichment scores (NESs) summarized enrichment direction: positive NESs indicated enrichment among genes positively associated with ASTS, whereas negative NESs indicated enrichment among genes negatively associated with ASTS. P values were adjusted using the Benjamini-Hochberg procedure^30^ within each collection and model.

Primary and canonical-driver-adjusted analyses were performed separately using the same procedure. Concordance between the two analyses was evaluated by Spearman rank correlation of NESs and by concordance of enrichment direction.

### Pharmacogenomic drug-response datasets and association analysis

PRISM Secondary dose-response curve parameters were obtained from secondary-screen-dose-response-curve-parameters.csv^31^. GDSC2 fitted dose-response data were obtained from GDSC2_fitted_dose_response_27Oct23.xlsx^3,32^ (release date, 27 October 2023). For PRISM, the response variable was log2-transformed Secondary AUC, with lower values indicating greater drug sensitivity. For GDSC2, the response variable was LN_IC50, for which lower values likewise indicate greater sensitivity. Drug-cell line records were matched to DepMap models for which ASTS scores and the required covariates were available. Compounds represented by fewer than 100 evaluable cell lines were excluded from both PRISM and GDSC2 analyses. A total of 108 compounds were represented in both datasets and were used for cross-screen comparison.

For each drug, a multivariable ordinary least-squares regression model was fitted with drug response as the dependent variable and ASTS as the variable of interest. Cancer lineage, recorded donor sex, and the proliferation proxy were included as baseline covariates. Heteroscedasticity-consistent HC3 standard errors^33^ were used for statistical inference. ASTS was z-standardized before model fitting, and the resulting ASTS coefficient was divided by the standard deviation of the corresponding drug-response variable to facilitate comparison among compounds and datasets. Positive standardized coefficients therefore indicate relative resistance in ASTS-high cells, whereas negative coefficients indicate relative sensitivity. Because lower response values indicate greater sensitivity in both PRISM and GDSC2, coefficient directions were directly comparable between the two screens.

### Cross-screen comparison and drug-class analysis

Compounds present in both PRISM and GDSC2 were matched using harmonized compound identifiers. Screen-specific ASTS effect estimates were compared by Spearman rank correlation and by concordance of effect direction. For the 108 compounds represented in both screens, ASTS effect estimates were converted to within-screen z scores by centering and scaling the drug-level coefficients across the matched compounds. For each drug, the cross-screen consensus score was calculated as the mean of its PRISM and GDSC2 z scores. Positive consensus values indicate associations shifted toward resistance relative to the screen-specific means of the matched compounds, whereas negative values indicate associations shifted toward sensitivity relative to those means. Sensitivity or resistance in ASTS-high cells was determined from the signs of the original screen-specific ASTS coefficients, not from the sign of the centered consensus score alone.

Drugs were grouped into mechanistic classes using annotated targets and pathway information. The principal classes examined included MEK-targeted drugs, EGFR-targeted drugs, EGFR-pathway drugs, ERK/MAPK-pathway drugs, PI3K/mTOR-pathway drugs, receptor tyrosine kinase (RTK)-pathway drugs, and non-EGFR RTK drugs. Class-level summaries were calculated from the consensus scores of the member drugs. In the harmonized GDSC2 annotations, the EGFR-target class was defined by EGFR in the putative-target field, whereas the EGFR-pathway class was defined by the EGFR-signaling pathway annotation. The EGFR-pathway class comprised gefitinib, erlotinib, lapatinib, osimertinib, and afatinib; the EGFR-target class additionally included foretinib. These were annotation-based groups and did not imply exclusive target selectivity.

### Adjustment for canonical oncogenic driver alterations

To evaluate whether ASTS-drug response associations were attributable to recurrent oncogenic driver alterations, hotspot mutation data were integrated with the ASTS metadata. Hotspot mutation calls were obtained from the DepMap Public 26Q1 hotspot mutation matrix^20^. Three composite binary driver variables were defined: MAPK-driver status from alterations in *KRAS*, *NRAS*, or *BRAF*; ERBB-driver status from alterations in *EGFR* or *ERBB2*; and PI3K-driver status from alterations in *PIK3CA*. Models with complete mutation information were included in this analysis.

Drug-response models were then refitted after addition of the MAPK-, ERBB-, and PI3K-driver variables to the baseline cancer lineage, donor sex, and proliferation covariates. The same adjustment strategy was applied separately to PRISM and GDSC2. Cross-screen effect concordance and drug-class summaries were recalculated from the driver-adjusted coefficients.

This analysis addressed the contribution of the specified canonical hotspot alterations but did not comprehensively account for copy-number changes, gene fusions, epigenetic alterations, or other genomic determinants.

### Cell-cycle and MYC adjustment

Additional sensitivity analyses evaluated the influence of cell-cycle- and MYC-related transcriptional programs on ASTS associations. Hallmark E2F targets, G2M checkpoint, and MYC targets V1 gene sets were used to generate transcriptional scores. Genes contained in the 78-gene ASTS signature and genes used in the simple proliferation proxy were removed before score calculation. After these exclusions, the E2F, G2M, E2F/G2M union, and MYC V1 sets contained 190, 192, 316, and 193 genes, respectively.

For each cell line, expression values of all available protein-coding genes were converted to within-sample percentile ranks using average ranks for tied values. The E2F, G2M, combined cell-cycle, and MYC V1 scores were calculated as the mean percentile rank of the genes belonging to the corresponding gene set. The combined cell-cycle score was calculated from the union of the E2F target and G2M checkpoint gene sets.

Three nested drug-response models were compared: the driver-adjusted model; the driver-adjusted model additionally including the combined cell-cycle score; and the driver-adjusted model additionally including both the cell-cycle and MYC V1 scores. Cross-screen concordance and drug-class effects were recalculated for each model. Cell-cycle adjustment was treated as a stringent sensitivity analysis because cell-cycle state could represent a biological component or mediator of ASTS rather than solely a confounder. The model additionally including both cell-cycle and MYC V1 scores was used for the individual EGFR/ERBB-targeted drug estimates in Fig. 3B.

### CRISPR dependency analysis

Gene dependency data were obtained from CRISPRGeneEffect.csv in DepMap release 26Q1^20^. More negative gene effect scores indicate stronger dependency; consequently, a positive ASTS regression coefficient indicates weaker dependency in ASTS-high cells, whereas a negative coefficient indicates stronger dependency. The released gene-effect scores were inferred using Chronos^34^.

Initial targeted analyses focused on *EGFR*, *ERBB2*, *MAP2K1*, and *MAP2K2*. Three nested models were examined: a baseline model including ASTS, cancer lineage, donor sex, and the proliferation proxy; a driver-adjusted model additionally including the MAPK-, ERBB-, and PI3K-driver variables; and a stringent model further including the cell-cycle and MYC V1 scores. Heteroscedasticity-consistent HC3 standard errors were used. In the stringent model, 1,135 cell lines were available for the targeted analysis. Stringent-model P values were corrected across the 26 genes in the analyzed targeted panel.

Composite dependency axes were constructed for the ERBB-direct pair (*EGFR* and *ERBB2*) and the MEK-direct pair (*MAP2K1* and *MAP2K2*). For each constituent gene, CRISPR gene effect values were z-standardized across cell lines. The composite score for each cell line was calculated as the arithmetic mean of the standardized gene effect values of the constituent genes, requiring nonmissing values for all genes in the corresponding axis. The same nested regression models used for the individual genes were then applied to the composite scores. Because higher CRISPR gene effect values represent weaker dependency, a positive ASTS coefficient for a composite axis indicates reduced dependency in ASTS-high cells. An exploratory ERBB-family composite was constructed analogously from *EGFR*, *ERBB2*, *ERBB3*, and *ERBB4*. Stringent-model P values were corrected across all three composite axes, separately from the individual-gene panel.

### Genome-wide CRISPR association analysis

Genome-wide CRISPR association analysis was performed using the stringent covariate model. For each gene, gene effect was modeled as a function of ASTS, cancer lineage, donor sex, the proliferation proxy, the MAPK-, ERBB-, and PI3K-driver variables, cell-cycle score, and MYC V1 score.

A total of 1,135 cell lines and 17,787 genes were included. For each gene, the ASTS coefficient, conventional model-based standard error, t statistic, and P value were obtained, and P values were corrected across the 17,787 genes using the Benjamini-Hochberg procedure. Positive coefficients identify genes for which ASTS-high cells show weaker dependency, whereas negative coefficients identify genes for which ASTS-high cells show stronger dependency. The same ranked genome-wide association statistics were used for pathway-level analyses.

### Adjustment for global CRISPR dependency burden

To test whether the mitochondrial signal reflected a generalized shift in CRISPR essentiality, two additional sample-level dependency covariates were constructed. Genes belonging to any of the mitochondrial gene sets used in the specificity analysis were excluded from both covariates. For the common-essential reference, eligible genes were required to have at least 95% coverage across cell lines and gene effect values below −0.5 in at least 80% of cell lines. The 500 eligible genes with the lowest median gene effect values were selected. The global common-essential burden score was the median gene effect value across these 500 genes within each cell line.

The second covariate, the gene-centered global dependency-shift score, was calculated across non-mitochondrial genes. For each gene, the median gene effect value across cell lines was subtracted from each cell line’s value. The score for each cell line was the median of these gene-centered values across non-mitochondrial genes.

Genome-wide gene effect models were then refitted after addition of both global dependency covariates to the stringent ASTS model. For interpretation of the mitochondrial phenotype, negative ASTS coefficients were taken to indicate increased dependency in ASTS-high cells. This analysis evaluates relative genetic dependency and does not imply increased mitochondrial transcription, oxidative phosphorylation rate, or oxygen consumption in ASTS-high cells.

### Mitochondrial gene-set enrichment and matched-gene permutation analysis

Mitochondrial pathway analyses used selected Hallmark and Reactome gene sets. The three primary sets were Hallmark oxidative phosphorylation, Reactome mitochondrial translation, and Reactome respiratory electron transport. The mitochondrial core union was defined as the union of these three sets.

Genes were ranked using the ASTS association statistics from the burden-adjusted genome-wide CRISPR models. To determine whether mitochondrial enrichment could be explained by the generally high baseline essentiality or distinctive variability of mitochondrial genes, a matched-gene permutation analysis was performed. Eligible control genes excluded all genes in the mitochondrial sets used in the specificity analysis and the 500 common-essential reference genes and were required to have a finite burden-adjusted ASTS t statistic and a finite, positive gene effect standard deviation. For matching, the median CRISPR gene effect value across cell lines and the natural logarithm of the gene effect standard deviation were calculated for each gene. Each matching feature was z-standardized across the eligible target and control genes.

For each mitochondrial target gene, squared Euclidean distance was calculated in this two-dimensional matching space, and the 100 nearest control genes were retained as its candidate matching pool. For each permutation, one matched control gene was sampled for every target gene without replacement within each random gene set. If no unused gene remained in a target-specific candidate pool, sampling was performed from the remaining eligible control genes. This procedure was repeated 10,000 times using a fixed random seed.

For each mitochondrial gene set, the observed mean burden-adjusted ASTS t statistic was compared with the distribution of mean t statistics from the matched random gene sets. A matched-permutation z score was calculated from the observed mean and the mean and standard deviation of the null distribution. The primary empirical P value tested whether the mitochondrial set was more negatively associated with ASTS than the matched random sets and was calculated using a plus-one correction^35^. P values were adjusted across the four tested mitochondrial gene sets using the Benjamini-Hochberg procedure. The analysis included 194 genes for oxidative phosphorylation, 100 for mitochondrial translation, 145 for respiratory electron transport, and 359 for the mitochondrial core union. With 10,000 permutations and the plus-one correction, the minimum attainable empirical P value was 1/10,001.

### Mechanism-guided pharmacological analysis of the mitochondrial dependency

Following identification of the mitochondrial genetic dependency, a mechanism-defined mitochondrial drug set was specified before examination of the corresponding drug-effect estimates. This analysis was therefore considered a post hoc, mechanism-guided pharmacological follow-up rather than a prospective validation.

The Tier 1 direct or near-direct oxidative phosphorylation (OXPHOS) set comprised IACS-010759^36^, BAY-87-2243^37^, rotenone, piericidin A^38^, antimycin A^39^, atovaquone^40^, and oligomycin A^41^. The Tier 2 broader mitochondrial set comprised phenformin, metformin^38^, elesclomol^42^, VLX600^43^, and devimistat^44^. Compound names were normalized and matched exactly to avoid substring-based false matches.

Available candidates were analyzed separately in each pharmacogenomic screen using the fully adjusted drug-response model containing cancer lineage, donor sex, the proliferation proxy, canonical driver variables, cell-cycle score, and MYC V1 score. Compounds represented by fewer than 100 evaluable cell lines were excluded. Heteroscedasticity-consistent HC3 standard errors were used for statistical inference. In PRISM, BAY-87-2243, oligomycin A, elesclomol, and VLX600 were evaluable; in GDSC2, elesclomol was evaluable.

CTRPv2 was used as an additional pharmacological resource^4,45^ for targeted follow-up of mitochondrial drug sensitivity. Drug-response data were extracted using PharmacoGx version 3.14.0 in R version 4.5.3^46^ from the CTRPv2 PharmacoSet obtained through ORCESTRA^47^ (PSet_CTRPv2.rds). After loading the original PharmacoSet, its object representation was updated using updateObject before extraction. The selected sensitivity measure was recomputed area above the curve (AAC; aac_recomputed). Because higher AAC values in the original metric indicate greater drug sensitivity, values were multiplied by −1 to generate a harmonized response for which lower values consistently indicated greater sensitivity. No further transformation was applied in the downstream ASTS association analysis. Duplicate records for the same compound and DepMap ModelID were aggregated using the median response. Oligomycin A was evaluable in 670 CTRPv2 cell lines and was analyzed using the corresponding fully adjusted regression model.

Candidate-level P values were adjusted using the Benjamini-Hochberg procedure both across all evaluable drugs within each screen and within the available mechanism-defined candidates in that screen. The candidate-level correction families contained four compounds in PRISM, one in GDSC2, and one in CTRPv2. Mechanism-level enrichment was additionally evaluated using matched-drug permutation. To account for differences in statistical precision and response scale among drugs, the number of evaluable cell lines and the standard deviation of the drug-response variable were each log1p-transformed and z-standardized across drugs. For each mitochondrial candidate drug, squared Euclidean distance was calculated in this two-dimensional matching space, and the 50 nearest noncandidate drugs were retained as its matching pool.

For each permutation, one matched background drug was sampled for every candidate drug without replacement within the random drug set. If no unused drug remained in a candidate-specific matching pool, sampling was performed from the remaining eligible background drugs. This procedure was repeated 10,000 times using fixed random seeds. For each candidate set, the observed mean standardized ASTS drug-response coefficient was compared with the distribution of mean coefficients from the matched random drug sets. A matched-permutation z score was calculated by subtracting the matched null mean from the observed mean and dividing by the matched null standard deviation. The empirical one-sided P value tested whether the candidate-set mean coefficient was more negative than those from matched random drug sets and was calculated using a plus-one correction. A negative standardized ASTS coefficient indicates greater drug sensitivity in ASTS-high cells. A negative matched-permutation z score indicates that the candidate-set mean coefficient is more negative than the matched null mean; its sign alone does not establish a negative ASTS coefficient. For drug-set permutation tests, Benjamini-Hochberg correction was applied jointly across the three evaluable PRISM/GDSC2 sets (two PRISM sets and one GDSC2 set), and separately across the two CTRPv2 sets. Both CTRPv2 sets contained only oligomycin A and represented the same compound-level comparison.

Because relatively few of the mechanism-defined mitochondrial compounds were available in each pharmacogenomic resource, these analyses were interpreted as pharmacological support for the mitochondrial genetic dependency rather than evidence of broad sensitivity to all mitochondrial or OXPHOS inhibitors.

### Leave-one-lineage-out analysis

The influence of individual cancer lineages was evaluated using leave-one-lineage-out (LOLO) analyses. Four primary functional endpoints were defined for this transportability audit: *EGFR* CRISPR dependency, a mitochondrial core CRISPR dependency composite, BAY-87-2243 response in PRISM, and oligomycin A response in PRISM. The five EGFR inhibitors analyzed in both PRISM and GDSC2 were evaluated as secondary pharmacological endpoints.

For each endpoint, lineage categories represented by at least 30 models in the endpoint dataset were eligible for sequential exclusion. The full model and each refitted model retained the other lineage categories, including those below this threshold. The model included ASTS, cancer lineage, donor sex, the proliferation proxy, the MAPK-, ERBB-, and PI3K-driver variables, cell-cycle score, and MYC V1 score. This model was repeatedly refitted after exclusion of one eligible lineage at a time. Each fitted model required at least 100 evaluable cell lines and at least 30 residual degrees of freedom. Heteroscedasticity-consistent HC3 standard errors were used throughout.

The minimum, maximum, and median standardized ASTS coefficients across lineage exclusions were recorded. Coefficients were direction-aligned: positive for EGFR dependency loss and EGFR-inhibitor resistance, and negative for mitochondrial dependency and mitochondrial-drug sensitivity. Positive aligned coefficients indicated the expected direction. Additional residual and influence diagnostics covered these 14 stringent endpoint models and the four Supplementary Fig. S3A score models. Endpoint sensitivity analyses excluded observations with leverage ≥ 1 − 10⁻⁸, with a further exclusion of the largest finite Cook’s-distance observation, or pooled lineages represented by fewer than five complete observations within each endpoint. Exclusion analyses retained the original standardization scales. Sensitivity P values were unadjusted and did not replace the primary tests or FDR values.

The mitochondrial core composite was constructed by z-standardizing each available core gene’s CRISPR gene effect values across cell lines and averaging the standardized values within each cell line. A score was retained only when at least 80% of the available core genes had nonmissing standardized values. This composite was a summary endpoint for influence assessment and did not replace the burden-adjusted mitochondrial pathway and matched-permutation analyses described above.

### Donor-sex-specific effects and interaction analyses

For donor sex analyses, cell lines annotated as Female or Male were retained, and those with unknown or missing sex annotations were excluded. Female- and male-derived cell lines were analyzed in a combined covariate-adjusted model containing ASTS, donor sex, an ASTS-by-donor-sex interaction, cancer lineage, the proliferation proxy, the MAPK-, ERBB-, and PI3K-driver variables, the cell-cycle score, and the MYC V1 score. Female was the reference category. Female-specific ASTS effects were obtained from the ASTS main coefficient; male-specific effects were obtained from the sum of the ASTS and interaction coefficients, with uncertainty calculated from the fitted covariance matrix. Heteroscedasticity-consistent HC3 standard errors were used. ASTS was standardized in the combined endpoint dataset, and both sex-specific effects were divided by the outcome standard deviation in that dataset.

Formal effect modification was assessed using the ASTS-by-donor-sex interaction term in the combined model. Male was coded as 1. The interaction coefficient therefore represented the male-minus-female difference in the ASTS slope. After alignment to the expected endpoint direction, a negative interaction coefficient indicated a larger effect in the expected direction in female-derived cell lines, whereas a positive coefficient indicated a larger effect in that direction in male-derived cell lines. Interaction P values were corrected jointly across all 14 tested endpoints using the Benjamini-Hochberg procedure.

Donor sex denotes the recorded sex of the individual from whom the cancer cell line was established. It was not used as a surrogate for androgen exposure, hormone concentration, or current androgen receptor signaling.

### Use of artificial intelligence

The authors used GPT-6.1sol (OpenAI) to assist with language revision, development of analysis scripts, and manuscript consistency checks. The authors reviewed and verified all AI-assisted content and take responsibility for the final manuscript.

### Statistical analysis

Continuous associations used Spearman correlation; multivariable analyses used linear regression. Conventional model-based standard errors were used for the genome-wide transcriptomic and CRISPR analyses, including burden-adjusted CRISPR models. Heteroscedasticity-consistent HC3 standard errors^33^ were used for drug-response models, targeted CRISPR analyses, and the LOLO and donor sex analyses. ASTS effect sizes were standardized for comparisons. Two-sided P values were used for primary regression tests. Multiple testing across genome-wide genes, drug panels, pathway analyses, or defined candidate sets was controlled using the Benjamini-Hochberg false discovery rate (FDR) procedure^30^, as appropriate to each analysis. Adjusted P values are reported as FDR values. FDR thresholds apply to each specified family, rather than globally across analyses. Linear regression estimated conditional associations for continuous outcomes. Fits used complete observations for required numerical and driver covariates, without imputation. Missing or unknown donor sex was retained as a category outside female–male interaction analyses. Lineages with fewer than five models in the metadata were pooled as OtherRare; missing lineage was encoded as Unknown or pooled. Displayed confidence intervals used coefficient ± 1.96 × SE. Residual-versus-fitted, normal Q-Q, and Cook’s-distance plots were inspected for the four Supplementary Fig. S3A score models.

Permutation analyses used empirical one-sided P values in the direction specified before each permutation test, with Benjamini-Hochberg FDR adjustment across the tested gene or drug sets. Statistical tests were not interpreted solely on the basis of nominal significance; effect direction, cross-screen concordance, covariate robustness, and sensitivity analyses were considered jointly.

## Results

### Global RNA-output suppression and relative transcriptional remodeling by DHT in zF cells

Our previous study demonstrated that DHT suppresses global gene expression in adrenal zona fasciculata (zF) cells. We first asked whether the DHT-associated reduction in global RNA output could be distinguished from gene-specific relative transcriptional remodeling within the same RNA-seq datasets. To this end, cell-number-normalized zF RNA-seq data were analyzed using two normalization strategies. ERCC-based normalization, which preserves differences in endogenous RNA abundance relative to the exogenous spike-in controls, revealed a broad negative shift in DHT-associated gene expression in both sexes. The median DHT-associated log2 fold changes were −0.235 in males and −0.646 in females. By contrast, standard DESeq2 normalization centered the distributions almost completely, with median log2 fold changes of 0.001 and 0.000 in males and females, respectively (Fig. 1A). Thus, ERCC-based normalization recapitulated the DHT-associated reduction in global RNA output, whereas standard normalization largely removed this global component and retained relative differences among genes. Concordance between independent counting procedures and additional normalization controls supported this interpretation (Fig. S1).

**Figure 1.**
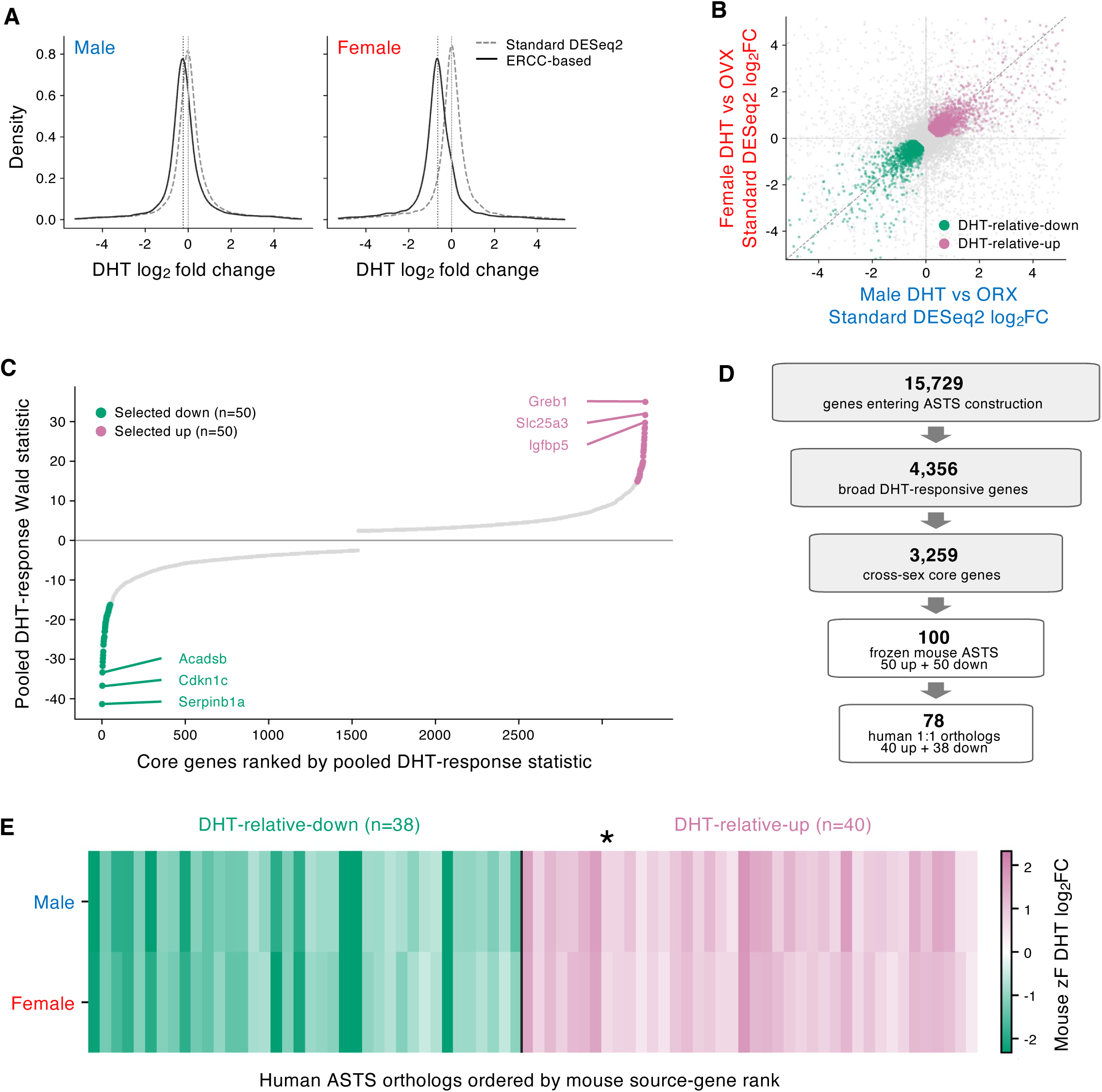
Construction of the androgen-suppressed transcriptional state signature from DHT-responsive genes in mouse adrenal zF cells. **(A)** Distributions of DHT-induced log2 fold changes in mouse adrenal zona fasciculata (zF) cells calculated using standard DESeq2 normalization (gray dashed lines) or ERCC-based normalization (black solid lines). Male and female zF cells were analyzed separately. A total of 16,535 genes were included. The median log2 fold changes obtained by standard and ERCC-based normalization were 0.001 and −0.235 in males and 0.000 and −0.646 in females, respectively. **(B)** Comparison of DHT-induced relative transcriptional responses between male and female zF cells after standard DESeq2 normalization. Each point represents one of the 15,729 genes entering ASTS construction. The Spearman correlation coefficient for all genes was ρ = 0.365. Genes included in the 3,259-gene cross-sex core are indicated as DHT-relative-up (magenta) or DHT-relative-down (green). The dashed diagonal indicates identity; horizontal and vertical reference lines indicate zero. **(C)** Ranking of the 3,259 cross-sex core genes according to the pooled DHT-response Wald statistic. The 50 genes with the strongest positive responses and the 50 genes with the strongest negative responses were selected to generate the 100-gene mouse ASTS signature. Representative genes are indicated. The signature was fixed before pharmacogenomic analyses were performed. **(D)** Flow diagram of ASTS construction. Among the 15,729 genes entering the analysis, 4,356 were classified as broadly DHT-responsive and 3,259 were retained as cross-sex core genes. Fifty DHT-relative-up and 50 DHT-relative-down genes were selected to generate the frozen 100-gene mouse ASTS signature. Orthology mapping identified 78 human one-to-one orthologs, consisting of 40 DHT-relative-up and 38 DHT-relative-down genes. **(E)** DHT-induced relative expression changes of the mouse source genes corresponding to the 78 human ASTS orthologs in male and female zF cells. Genes are ordered according to their direction-specific rank in the frozen mouse signature. The asterisk marks MGARP (mouse Mgarp), the lower-confidence ortholog excluded from the 77-gene sensitivity set. Green and magenta denote DHT-relative-down and DHT-relative-up genes, respectively. See also Figures S1 and S2.

We therefore used standard DESeq2 normalization to characterize relative transcriptional responses to DHT. Among the 15,729 genes entering signature construction, DHT effects in males and females were positively correlated (Spearman ρ = 0.365; Fig. 1B). Application of the criteria described in Methods identified 4,356 broadly DHT-responsive genes, of which 3,259 also met the effect-size thresholds and the criterion excluding strong sex-by-DHT interactions and were retained as the cross-sex core (Fig. 1D). These results indicate that, in addition to reducing global RNA output, DHT induces a relative transcriptional remodeling program that is shared, at least in part, between male and female zF cells.

To construct a compact signature representing this relative response, the 3,259 cross-sex core genes were ranked according to the pooled DHT-effect Wald statistic. The 50 highest-ranking genes in each direction were selected, yielding a 100-gene mouse signature comprising 50 DHT-relative-up and 50 DHT-relative-down genes (Fig. 1C,D). The signature was fixed before any pharmacogenomic analyses were performed. E2 responses of both the cross-sex core and the selected signature genes were subsequently examined as a specificity analysis and were not used for signature selection (Fig. S2A,B).

The frozen mouse signature was then projected to human genes by one-to-one orthology mapping. Seventy-eight genes satisfied the primary mapping criterion, comprising 40 DHT-relative-up and 38 DHT-relative-down genes (Fig. 1D and Fig. S2C,D). A 77-gene high-confidence sensitivity set was generated by excluding the single one-to-one ortholog with a lower Ensembl orthology-confidence annotation. The source genes retained concordant relative DHT responses between male and female zF cells (Fig. 1E). These 78 human orthologs were used to define the androgen-suppressed transcriptional state (ASTS) score in subsequent analyses. Importantly, ASTS represents resemblance to the relative DHT-associated transcriptional state and does not quantify absolute RNA output per cell; therefore, a higher ASTS score should not be interpreted as lower absolute transcriptional output.

### Construction of the ASTS score in human cancer cell lines

We next asked whether the zF-derived relative DHT-associated transcriptional state could be quantified in human cancer cells. The 78 human ASTS genes were projected onto the DepMap expression dataset, and genes were ranked within each cell line to minimize dependence on absolute expression scale. The ASTS score was calculated as the difference between the mean percentile ranks of the DHT-relative-up and DHT-relative-down genes. After restricting the analysis to the default expression profile for each DepMap model, ASTS scores were obtained for 1,719 unique cancer cell lines (Fig. 2A).

**Figure 2.**
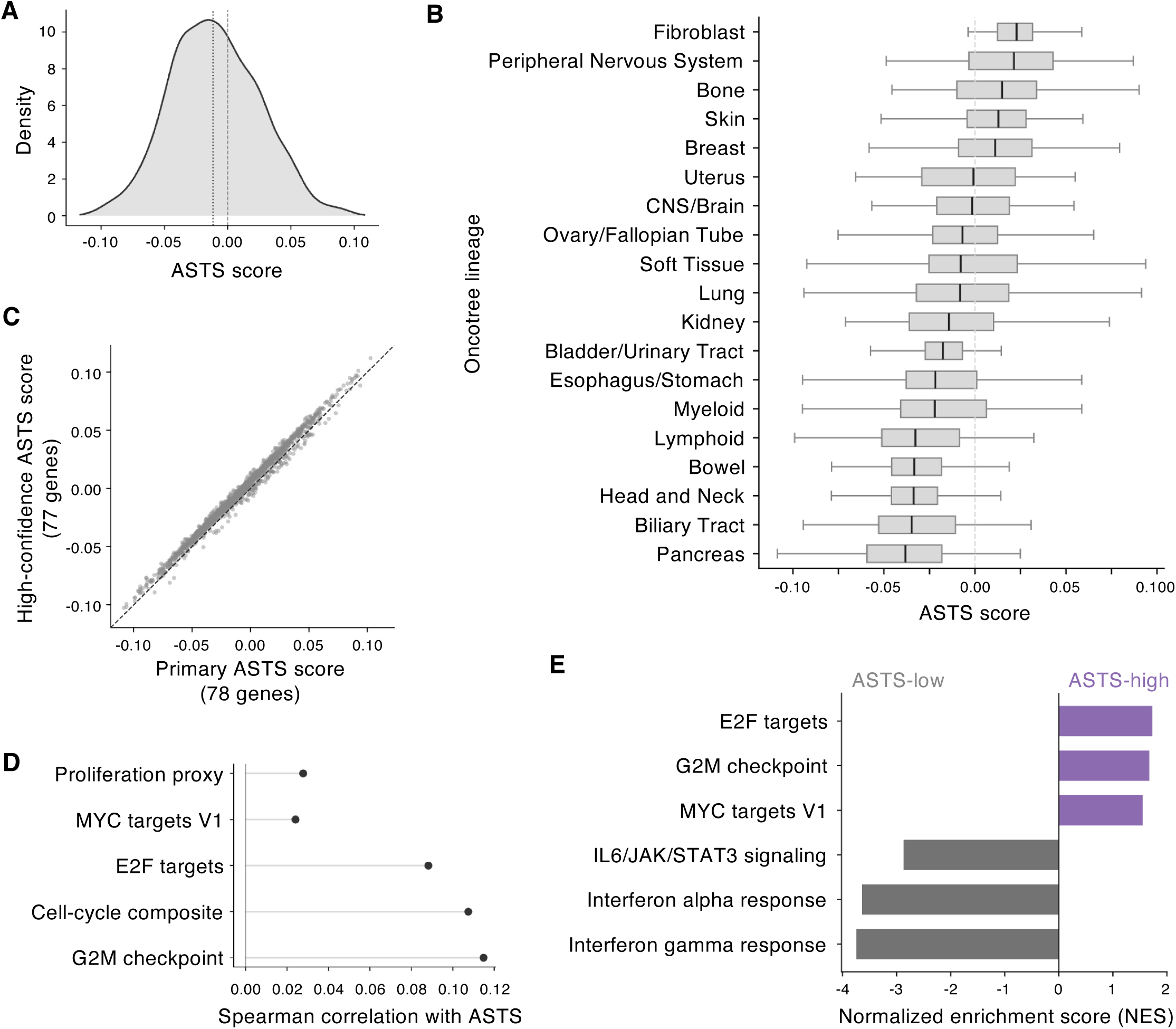
Distribution and transcriptional characteristics of ASTS across human cancer cell lines. **(A)** Distribution of the primary 78-gene ASTS score across 1,719 unique cancer cell lines in the DepMap expression dataset. ASTS was calculated from within-sample expression ranks of DHT-relative-up and DHT-relative-down genes. The dashed vertical line marks zero; the dotted line marks the median. **(B)** Distribution of ASTS scores across major OncoTree lineages. Boxplots show the median and interquartile range, with whiskers extending to 1.5 times the interquartile range. Lineage was included as a covariate in subsequent pharmacogenomic and CRISPR dependency analyses. Only lineages with at least 30 cell lines were shown (19 lineages; 1,550 cell lines in total); lineage-specific counts are provided in Source Data. **(C)** Comparison of ASTS scores calculated using the primary 78-gene signature and the 77-gene high-confidence ortholog sensitivity set. Each point represents one cell line. The two scores were highly correlated (Spearman ρ = 0.995). The dashed diagonal indicates identity. **(D)** Spearman correlations between ASTS and transcriptional scores representing proliferation, MYC targets V1, E2F targets, a composite cell-cycle score, and the G2M checkpoint. The correlation between ASTS and the simple proliferation proxy was ρ = 0.028. **(E)** Selected Hallmark gene sets associated with ASTS in the driver-adjusted genome-wide transcriptome analysis. Normalized enrichment scores (NESs) are shown. Positive values indicate enrichment in ASTS-high cells and negative values indicate enrichment in ASTS-low cells. Selected ASTS-high pathways include E2F targets, G2M checkpoint, and MYC targets V1, whereas ASTS-low pathways include IL6/JAK/STAT3 signaling and interferon responses. See also Figures S2 and S3. Purple and gray bars denote positive and negative NESs, respectively.

ASTS scores varied substantially among cancer lineages (Fig. 2B), indicating that lineage could contribute to apparent ASTS-associated phenotypes. Cancer lineage was therefore included as a categorical covariate in subsequent transcriptomic, pharmacogenomic, and CRISPR dependency analyses. Despite this lineage-associated variation, the ASTS score was highly robust to orthology-confidence filtering: scores calculated using the 78-gene primary signature and the 77-gene high-confidence sensitivity set were nearly identical (Spearman ρ = 0.995; Fig. 2C and Fig. S2E).

Because proliferation is a major determinant of both cancer-cell transcriptomes and drug responses^48^, we next examined whether ASTS primarily reflected proliferative state. ASTS showed essentially no correlation with the simple proliferation proxy (Spearman ρ = 0.028; Fig. 2D). By contrast, weak positive associations were observed with independently constructed E2F-, G2M-, and MYC-related transcriptional scores, and these associations remained detectable after multivariable adjustment (Fig. S3A). Thus, ASTS was not equivalent to a general proliferation index, although it was associated with specific cell-cycle-related transcriptional programs.

To further characterize the transcriptional architecture associated with ASTS, we performed genome-wide transcriptomic association analysis followed by gene-set enrichment analysis. In the canonical-driver-adjusted analysis, ASTS-high cells were enriched for E2F targets, G2M checkpoint, and MYC-related programs, whereas ASTS-low cells were enriched for interferon-response, inflammatory, and IL6/JAK/STAT3-related programs (Fig. 2E). The overall enrichment pattern was highly concordant between the primary and driver-adjusted analyses (Fig. S3B,C), indicating that the major transcriptional features associated with ASTS were not substantially altered by adjustment for the canonical oncogenic driver variables examined here.

These results characterize ASTS as a relative transcriptional score that is robust to orthology-confidence filtering and is associated with lineage and selected cell-cycle-related programs.

### Association of ASTS with EGFR inhibitor response and EGFR dependency

We next investigated whether ASTS was associated with drug response. Associations between ASTS and drug response estimated separately in PRISM and GDSC2 showed a positive cross-screen correlation (Spearman ρ = 0.391; Fig. 3A). Several inhibitors of the EGFR and MEK pathways had coefficients above the respective screen-specific means.

**Figure 3.**
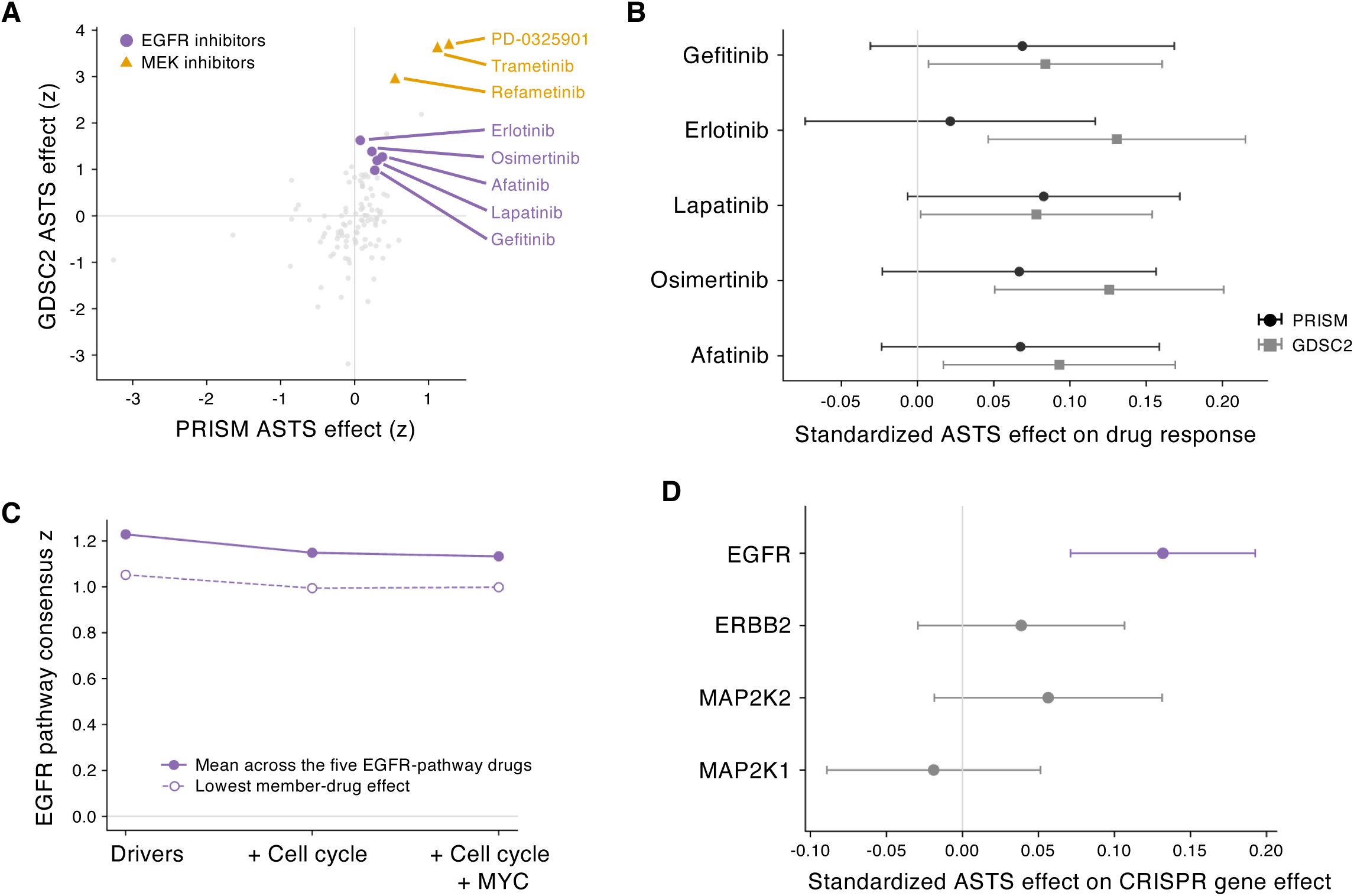
Higher ASTS is associated with reduced sensitivity to EGFR/ERBB-targeted drugs and lower EGFR dependency. **(A)** ASTS-associated drug-response effects in PRISM and GDSC2 from baseline models adjusted for lineage, donor sex, and proliferation. Each gray point represents a drug measured in both datasets. EGFR inhibitors are highlighted in purple and MEK inhibitors in orange. Values are coefficients centered and scaled across the 108 matched drugs within each screen. Positive z scores indicate effects above the corresponding screen-specific mean; resistance or sensitivity is determined by the original coefficient signs. Spearman ρ = 0.391. **(B)** ASTS-associated responses to gefitinib, erlotinib, lapatinib, osimertinib, and afatinib in PRISM and GDSC2 using fully adjusted models that additionally included canonical hotspot-driver variables, cell-cycle score, and MYC V1 score. Points indicate standardized ASTS regression coefficients and horizontal lines indicate 95% confidence intervals. Positive coefficients indicate reduced drug sensitivity in ASTS-high cells. All ten coefficients were positive; some confidence intervals crossed zero. Sample sizes (PRISM/GDSC2): gefitinib 461/699, erlotinib 473/696, lapatinib 473/705, osimertinib 473/698, and afatinib 465/705. Black circles: PRISM; gray squares: GDSC2. **(C)** Robustness of the EGFR-pathway drug-response association to sequential covariate adjustment. Filled circles indicate the mean centered consensus z score across the five EGFR-pathway drugs defined in Methods; open circles indicate the lowest member-drug consensus z score. Consensus scores are the mean of the two screen-specific centered z scores described in panel A. Models retained baseline covariates and additionally included driver variables, drivers plus cell-cycle score, or drivers plus cell-cycle and MYC V1 scores. **(D)** Association between ASTS and CRISPR dependency on EGFR, ERBB2, MAP2K2, and MAP2K1 using the stringent targeted CRISPR model with panel B covariates. Points indicate standardized regression coefficients and horizontal lines indicate 95% confidence intervals. More negative CRISPR gene effects indicate stronger dependency; positive ASTS coefficients indicate reduced ASTS-high dependency. EGFR showed a significant positive association (standardized β = 0.132, P = 0.0000209, FDR = 0.000543), whereas ERBB2, MAP2K1, and MAP2K2 did not reach significance. See also Figure S4. **FDR was corrected across the 26-gene panel.** Each gene included 1,135 cell lines. Purple denotes EGFR; gray denotes the other genes.

In the fully adjusted models, including lineage, donor sex, proliferation, canonical hotspot-driver variables, cell-cycle score, and MYC V1 score, all ten effect estimates for the five selected EGFR/ERBB-targeted drugs (gefitinib, erlotinib, lapatinib, osimertinib, and afatinib) were positive (Fig. 3B). Individual estimates did not uniformly reach statistical significance. Sequential adjustment analyses are summarized in Fig. 3C. The EGFR-pathway drug class retained a positive mean centered consensus z score after addition of MAPK-, ERBB-, and PI3K-driver covariates, with only modest changes after further adjustment for cell-cycle and MYC scores (Fig. 3C). Similar analyses of other drug classes are shown in Fig. S4A,B.

These pharmacological observations raised the possibility that ASTS-high cells have altered dependency on EGFR itself. We therefore analyzed DepMap CRISPR gene-effect data. In the stringent multivariable model, ASTS was positively associated with the EGFR gene-effect score (standardized β = 0.132, P = 0.0000209, 26-gene-panel FDR = 0.000543), indicating reduced EGFR dependency in ASTS-high cells (Fig. 3D). By contrast, the associations with ERBB2, MAP2K1, and MAP2K2 were weaker and did not reach significance.

The MEK inhibitor phenotype differed from the EGFR phenotype. Several MEK inhibitors also showed positive ASTS-associated resistance effects, whereas neither MAP2K1 nor MAP2K2 showed a corresponding reduction in genetic dependency. Consistent with this distinction, the ERBB-direct composite dependency was positively associated with ASTS, whereas the MAP2K1/MAP2K2 composite showed little evidence of reduced dependency (Fig. S4C,D). Taken together, these results indicate that ASTS-associated resistance to EGFR-targeted drugs is accompanied by reduced EGFR dependency, whereas the MEK inhibitor resistance phenotype cannot be explained simply by reduced MAP2K1/2 dependency.

### Selective mitochondrial dependency in ASTS-high cells

We next performed a genome-wide CRISPR analysis to identify genetic dependencies associated with ASTS without restricting the analysis to predefined signaling pathways. Among the 17,787 genes examined, multiple mitochondrial genes were located toward the ASTS-high-dependent end of the distribution, including SLC25A51, MTIF3, RMND1, NDUFB4, MRPS11, and LRPPRC (Fig. 4A).

**Figure 4.**
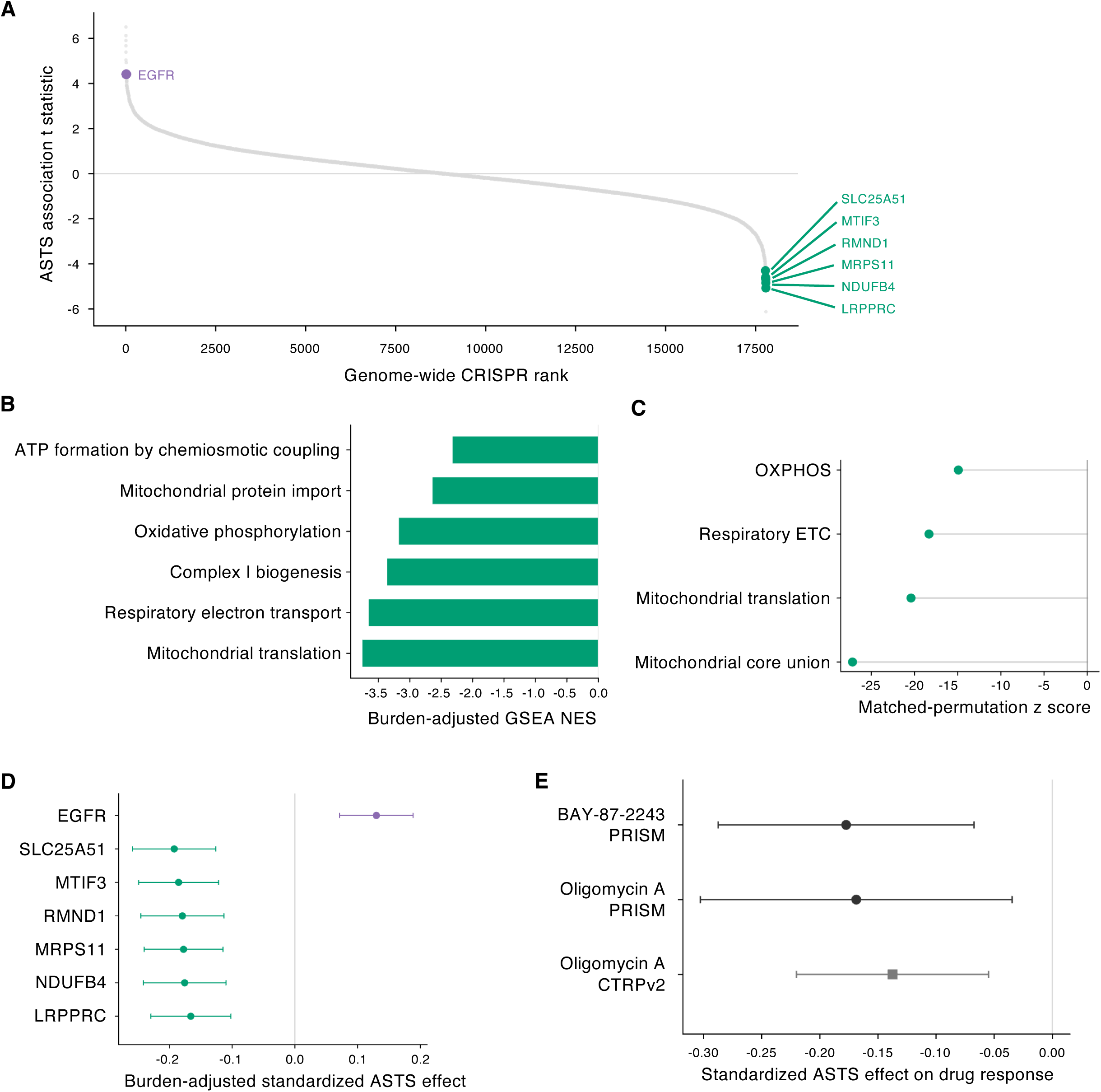
Higher ASTS is associated with selective mitochondrial genetic dependency and sensitivity to selected mitochondrial perturbagens. **(A)** Genome-wide ASTS–CRISPR associations across 17,787 genes, ranked by the ASTS t statistic. The stringent model included lineage, donor sex, proliferation, canonical hotspot-driver variables, cell-cycle score, and MYC V1 score. EGFR is purple; SLC25A51, MTIF3, RMND1, MRPS11, NDUFB4, and LRPPRC are green. Negative statistics indicate stronger dependency in ASTS-high cells. **(B)** Selected mitochondrial pathways from GSEA of the stringent model additionally adjusted for global common-essential burden and a gene-centered global dependency shift. Bars show NESs; negative values indicate stronger ASTS-high dependency. Pathways include chemiosmotic ATP formation, mitochondrial protein import, oxidative phosphorylation, complex I biogenesis, respiratory electron transport, and mitochondrial translation. **(C)** Matched-gene permutation analysis using panel B statistics. Negative matched-null z scores indicate stronger ASTS-high dependency than expected from genes matched for baseline essentiality and variability. Values were −14.9 (oxidative phosphorylation), −18.3 (respiratory electron transport), −20.4 (mitochondrial translation), and −27.2 (mitochondrial core union). All four sets had empirical one-sided P = 1/10,001 and four-set FDR = 0.000100. OXPHOS, oxidative phosphorylation; ETC, electron transport chain. **(D)** Burden-adjusted standardized ASTS coefficients using panel B covariates. EGFR is purple and mitochondrial genes are green. Negative coefficients indicate stronger ASTS-high dependency. Horizontal lines indicate 95% confidence intervals. Sample sizes were 1,092 cell lines for SLC25A51 and 1,135 for each other displayed gene. **(E)** Standardized ASTS drug-response coefficients with 95% confidence intervals; negative values indicate greater sensitivity. PRISM BAY-87-2243 (n = 457): β = −0.177, P = 0.00158, candidate-set FDR = 0.00634, screen-wide FDR = 0.222; PRISM oligomycin A (n = 360): β = −0.169, P = 0.0137, candidate-set FDR = 0.0273, screen-wide FDR = 0.469; CTRPv2 oligomycin A (n = 670): β = −0.137, P = 0.00112, candidate-set FDR = 0.00112, screen-wide FDR = 0.0305. These are exploratory pharmacological associations. See also Figure S5.

Because mitochondrial genes are frequently essential across many cancer cell lines, we next asked whether this pattern could reflect a generalized shift in CRISPR dependency. We therefore refitted the genome-wide models after addition of two global dependency covariates representing common-essential burden and a gene-centered global dependency shift. Gene-set enrichment analysis of these burden-adjusted associations revealed strong enrichment of mitochondrial pathways among genes showing increased dependency in ASTS-high cells. Oxidative phosphorylation showed a markedly negative enrichment score (NES = −3.17), with still stronger negative enrichment for mitochondrial translation and respiratory electron transport (NES = −3.76 and −3.65, respectively). Complex I biogenesis, mitochondrial protein import, and ATP formation by chemiosmotic coupling were enriched in the same direction (Fig. 4B).

We further compared predefined mitochondrial gene sets with control genes matched for baseline essentiality and gene-effect variability. The matched-permutation z scores were −14.9 for oxidative phosphorylation, −20.4 for mitochondrial translation, −18.3 for respiratory electron transport, and −27.2 for the mitochondrial core union (Fig. 4C). Thus, the mitochondrial dependency was substantially stronger than expected from the baseline essentiality and variability of the constituent genes. The relationships between ASTS and the two global CRISPR burden metrics are shown in Fig. S5A,B. Consistent with these pathway-level analyses, individual mitochondrial genes, including SLC25A51, MTIF3, RMND1, MRPS11, NDUFB4, and LRPPRC, retained negative ASTS associations after adjustment for global dependency burden, whereas EGFR retained the opposite positive association (Fig. 4D). All four matched-gene sets had an empirical one-sided P value of 1/10,001 and a four-set FDR of 0.000100.

We next asked whether this mitochondrial genetic dependency was accompanied by pharmacological sensitivity to mitochondrial perturbation. As a post hoc, mechanism-guided analysis, we evaluated a mechanism-defined set of mitochondrial perturbagens specified before examination of the corresponding drug-effect estimates. In PRISM, ASTS-high cells showed greater sensitivity to BAY-87-2243 (standardized β = −0.177) and oligomycin A (standardized β = −0.169). Oligomycin A showed a concordant negative association in CTRPv2 (standardized β = −0.137; Fig. 4E). The PRISM associations met FDR < 0.05 within the mechanism-defined candidate set but not across all evaluable PRISM drugs. The CTRPv2 oligomycin A association met both candidate-set and screen-wide FDR < 0.05. These findings were treated as exploratory pharmacological support. At the mechanism level, the direct OXPHOS inhibitor set in PRISM was shifted toward more negative ASTS drug-response effects relative to matched background drugs. The availability and responses of the complete mitochondrial candidate set, together with matched-drug permutation analyses, are shown in Fig. S5C–E. Together, these results indicate that ASTS-high cells exhibit a selective mitochondrial genetic dependency, with concordant pharmacological sensitivity to mitochondrial perturbation in the compounds and screens available for analysis.

### Robustness of ASTS-associated functional phenotypes across cell lineages and donor sex

Because ASTS scores differed among cancer lineages, we finally examined whether the major functional associations described above were driven by particular lineages. Leave-one-lineage-out analyses were performed for four primary endpoints: EGFR dependency, mitochondrial-core dependency, and sensitivity to BAY-87-2243 and oligomycin A. Exclusion of each of 18 major lineages preserved the expected direction of the EGFR and mitochondrial-core CRISPR associations. The BAY-87-2243 and oligomycin A associations likewise retained their expected directions across all evaluable lineage exclusions, comprising five and three major lineages, respectively (Fig. 5A).

**Figure 5.**
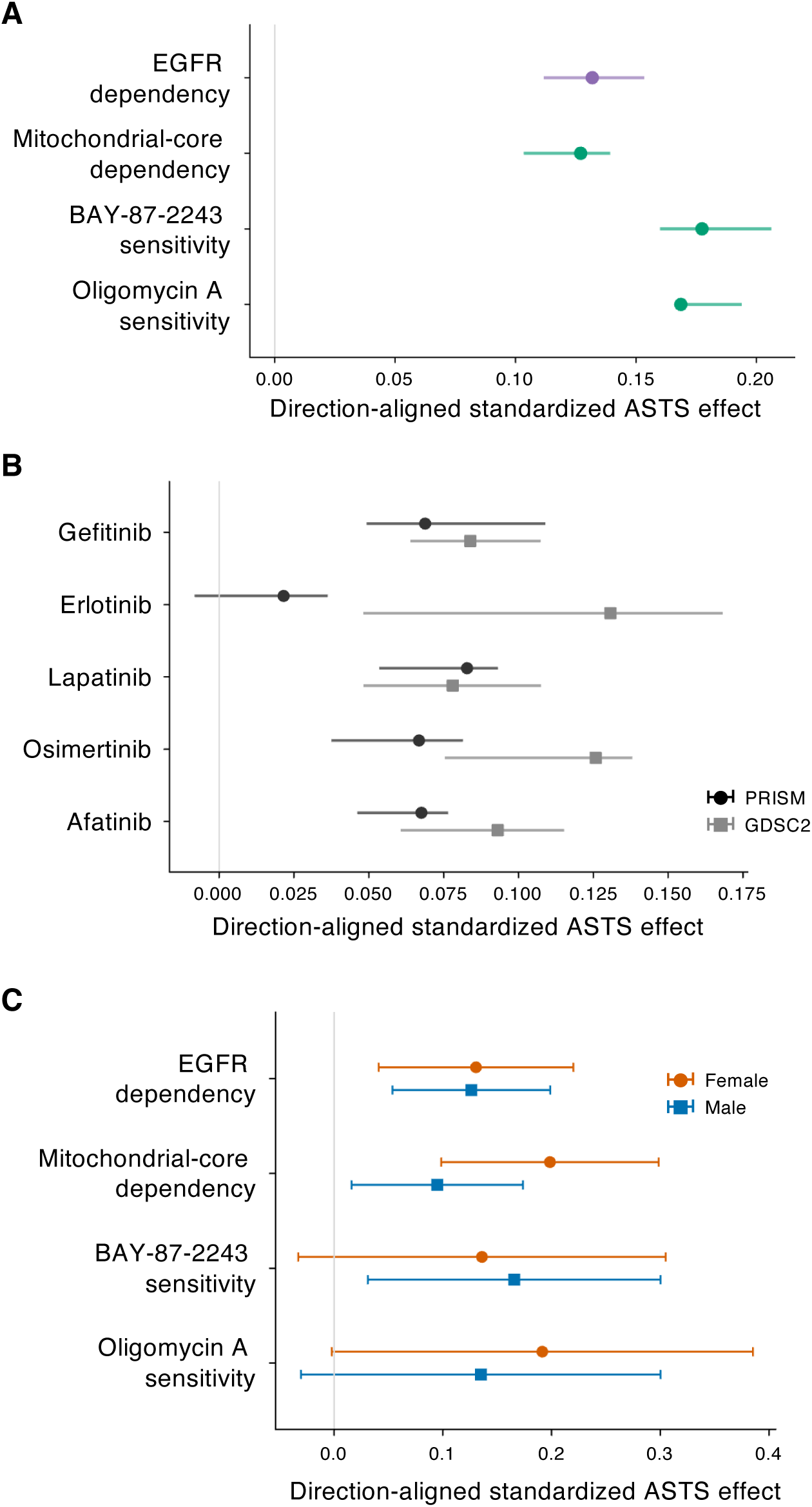
Robustness of major ASTS-associated functional phenotypes across cancer lineages and cell-line donor sex. **(A)** Leave-one-lineage-out analysis of four primary functional endpoints defined for the transportability audit. Points indicate the standardized ASTS effect obtained from the full model, and horizontal lines indicate the minimum-to-maximum range obtained after exclusion of individual major cancer lineages. Effects were direction-aligned so that positive values support the predefined phenotype. EGFR dependency and mitochondrial-core dependency were evaluated across 18 lineage exclusions; BAY-87-2243 and oligomycin A sensitivity were evaluable across five and three major lineage exclusions, respectively. All four endpoints retained the predefined direction across all tested lineage exclusions. Full and refitted models included lineage, donor sex, proliferation, canonical hotspot-driver variables, cell-cycle score, and MYC V1 score. The mitochondrial-core composite models did not include the global CRISPR dependency-burden covariates used in Fig. 4B-D. **(B)** Leave-one-lineage-out analysis of ASTS-associated resistance to five EGFR inhibitors in PRISM and GDSC2. Points indicate full-model standardized effects, and horizontal lines indicate the range across lineage exclusions. Effects were direction-aligned so that positive values indicate ASTS-high resistance. Nine of the ten drug-response estimates retained the expected direction across all lineage exclusions; the PRISM erlotinib association showed one estimate slightly crossing zero. **(C)** Sex-specific ASTS slopes estimated from the combined ASTS-by-donor-sex interaction model are shown for the four primary endpoints. Female-derived cell lines are shown in orange and male-derived cell lines in blue. Points indicate direction-aligned standardized ASTS coefficients and horizontal lines indicate 95% confidence intervals. For EGFR and mitochondrial-core dependency, 1,082 cell lines were analyzed (465 female and 617 male); for BAY-87-2243, 438 cell lines were analyzed (190 female and 248 male); and for oligomycin A, 342 cell lines were analyzed (151 female and 191 male). Formal ASTS-by-donor-sex interaction analyses are shown in Figure S6. Donor sex indicates the recorded sex of the individual from whom the cell line was derived and does not represent androgen exposure. See also Figure S6.

We next extended this analysis to the five EGFR inhibitors evaluated in both PRISM and GDSC2. Nine of the ten drug-response estimates retained the expected direction across all tested lineage exclusions (Fig. 5B). The exception was erlotinib in PRISM, for which one leave-one-lineage-out estimate crossed slightly into the opposite direction. Thus, although the degree of robustness varied among individual drugs, the overall EGFR inhibitor phenotype was not attributable to a single major cancer lineage. Additional endpoint diagnostics identified unit-leverage observations from singleton lineages. Their exclusion had negligible effects on ASTS coefficients and HC3 standard errors. Pooling lineages with fewer than five observations within each endpoint preserved coefficient signs for all 14 models. The four primary endpoints retained their expected directions and unadjusted P < 0.05 in these sensitivity analyses, including further removal of the largest finite Cook’s-distance observation. Individual drug significance was less stable: the GDSC2 gefitinib association changed from unadjusted P = 0.032 to P = 0.051 after that exclusion.

Finally, we examined whether the four primary functional associations differed according to the recorded donor sex of the cell lines. The direction-aligned ASTS effects were directionally concordant between female- and male-derived cell lines for all four endpoints (Fig. 5C). Formal ASTS-by-donor-sex interaction analyses provided no statistically supported evidence of effect modification after correction for multiple testing, either for the four primary endpoints or for the EGFR inhibitor responses (Fig. S6). Thus, the available data provide no strong evidence that recorded donor sex substantially modifies the major ASTS-associated functional phenotypes examined here, although these analyses do not establish equivalence between sexes. Importantly, donor sex denotes the recorded sex of the individual from whom the cell line was established and should not be interpreted as a surrogate for androgen exposure or current androgen receptor signaling.

## Discussion

This study examined whether a transcriptional state derived from a defined in vivo hormonal perturbation could remain biologically informative when transferred across species and cellular contexts. ASTS was constructed from relative DHT-associated transcriptional remodeling in isolated mouse adrenal zona fasciculata cells and was fixed before interrogation of cancer pharmacogenomic and dependency data. The term ASTS therefore denotes resemblance to the relative transcriptional state associated with DHT-mediated transcriptional suppression in zF cells, rather than a direct measure of androgen exposure, androgen receptor activity, or absolute RNA output in cancer cells. Within these boundaries, the association of ASTS with pharmacological and genetic phenotypes measured in separate screens suggests that the transcriptional information captured by the score is not restricted to the tissue in which it was originally defined.

Gene-expression signatures have long provided a means of transferring biological information across experimental contexts. The Connectivity Map established that transcriptional responses can reveal functional relationships among drugs, genetic perturbations, physiological processes, and disease states, and the L1000 framework subsequently extended this principle to perturbational profiling at much larger scale^10,11^. More recently, integration of transcriptional states with genome-scale dependency data has shown that cancer cell states can expose selective genetic vulnerabilities; for example, Killarney et al. identified PKN2 as a dependency associated with the mesenchymal-like cancer cell state across diverse lineages^8^. ASTS extends this general concept in a different direction because its defining information originates not from a cancer phenotype or treatment-resistance program, but from the transcriptional response of nonmalignant adrenal cells to an in vivo endocrine perturbation. We therefore view ASTS primarily as a state descriptor for hypothesis generation and functional interrogation, rather than as a validated predictive biomarker. The present study provides a proof of concept that a transcriptional state defined in one biological context can remain informative when projected into a distinct system and integrated with separately generated functional-genomic resources. Because the associated functional phenotypes remained evident after accounting for canonical oncogenic driver alterations, transcriptional state information may complement genotype-based descriptions of cancer when prioritizing candidate therapeutic vulnerabilities. Related work has integrated transcriptional states, CRISPR dependencies, and drug responses in head and neck cancer, including ex vivo evaluation in patient-derived three-dimensional microtumors^49^.

One potentially important feature of the ASTS-associated phenotype is the coexistence of altered dependence on growth-factor signaling and increased mitochondrial dependency. Previous studies provide several precedents for an association between escape from dominant oncogenic signaling and reliance on mitochondrial function. In EGFR-mutant lung adenocarcinoma, osimertinib treatment suppresses glycolysis and creates a strong requirement for mitochondrial oxidative phosphorylation, whereas inhibition of mitochondrial respiration delays the emergence of resistance^50^. Mitochondrial adaptations have also been identified in treatment-resistant states driven by MAPK pathway inhibition. A subset of melanomas resistant to MEK inhibition exhibits PGC1α-associated oxidative metabolism, whereas trametinib-resistant KRAS-mutant lung cancer cells can increase mitochondrial oxidative metabolism through enhanced pyruvate and fatty-acid utilization^51,52^. Experiments involving KRAS ablation in pancreatic cancer similarly showed that residual cells surviving loss of KRAS-driven signaling become highly dependent on mitochondrial function^53^. These systems differ in tumor type, treatment history, and mechanism, and mitochondrial reliance is clearly not a universal consequence of targeted-therapy resistance. Nevertheless, they establish that reduced reliance on a dominant signaling axis and increased mitochondrial requirements can coexist in defined cancer cell states. Our findings raise the possibility that ASTS identifies another cellular context in which these properties coexist. They do not establish that reduced signaling dependence causes mitochondrial dependency, that mitochondrial dependency causes drug resistance, or that either phenotype precedes the other. Expression-based EMT states have also been associated with EGFR-inhibitor resistance in non-small-cell lung cancer^54^; the present analyses do not establish that ASTS is independent of EMT. More recent preclinical work links PGC1β-dependent mitochondrial biogenesis to osimertinib resistance^55^.

The mitochondrial phenotype associated with ASTS should also be distinguished from an OXPHOS-high metabolic state. A genetic requirement for mitochondrial pathways does not establish increased respiratory flux, oxygen consumption, or mitochondrial ATP production. Mitochondrial translation represents one particularly interesting component of this phenotype. MYC is known to coordinate the expression of numerous nuclear-encoded mitochondrial genes and to promote mitochondrial biogenesis, and MYC-driven lymphomas can acquire a selective requirement for mitochondrial translation machinery^56,57^. These observations provide a plausible biological connection to the MYC-related transcriptional features associated with ASTS, but they are unlikely to provide a complete explanation because the mitochondrial dependency remained evident after accounting for MYC- and cell-cycle-related transcriptional programs. Moreover, mitochondrial respiration supports proliferating cells through functions beyond ATP production. Independent studies showed that electron transport can be essential for maintaining the redox conditions required for aspartate biosynthesis and thereby support anabolic proliferation^58,59^. The mitochondrial phenotype associated with ASTS could therefore reflect requirements for mitochondrial translation, redox homeostasis, biosynthetic metabolism, bioenergetic function, or combinations of these processes. It is consequently more appropriately interpreted as a functional mitochondrial dependency than as evidence of increased oxidative phosphorylation activity.

The adrenal origin of ASTS provides a biological context for considering the relationship between transcriptional state and metabolism. In steroidogenic cells, Ad4BP/SF-1 (NR5A1) coordinates glycolytic and cholesterol-biosynthetic programs^13,14^, and the DHT-responsive zF system similarly links androgen signaling, NR5A1 expression, transcriptional output, and cellular metabolism^12^. NR5A1 dosage is also relevant to adrenocortical neoplasia. Increased copy number and expression have been reported particularly in childhood adrenocortical tumors, experimentally increased SF-1 dosage promotes adrenocortical cell proliferation and tumor formation, and high SF-1 expression in adult adrenocortical carcinoma has been associated with adverse outcome^60–63^. These observations underscore the dosage sensitivity of the adrenal regulatory circuitry from which ASTS originated. They do not, however, imply that ASTS is an NR5A1-activity signature or that the regulatory mechanism linking DHT exposure, androgen receptor signaling, and NR5A1 expression in normal zF cells is preserved across the cancer cell lines analyzed here. Rather, DHT-responsive remodeling may capture part of a broader configuration linking transcriptional regulation and metabolism that remains informative outside its original adrenal context.

Several limitations define the scope of these conclusions. First, ASTS was transferred from mouse adrenal cells to human cancer cells through one-to-one orthology, which necessarily preserves only part of the regulatory information in the original state and cannot reproduce species- or tissue-specific transcriptional circuitry. Second, the present analyses do not establish that ASTS is uniquely informative relative to other possible transcriptional signatures, nor do they demonstrate that experimentally altering the ASTS-associated state will change drug response or genetic dependency. ASTS should therefore be regarded as a useful state representation identified in the present analysis rather than as a uniquely optimized classifier. Third, the mitochondrial phenotype has not yet been resolved by direct measurements of respiration, metabolite flux, mitochondrial translation, or biosynthesis, and pharmacological coverage of mitochondrial perturbagens was limited. The genetic dependency should therefore not be extrapolated to broad sensitivity to all mitochondrial or OXPHOS inhibitors. Fourth, the analyses were performed predominantly in established cancer cell lines and do not establish clinical predictive utility in tumors or patients. The pharmacogenomic screens and CRISPR data were generated separately but include overlapping cell-line models and share DepMap-derived ASTS scores and covariates. Their concordance provides cross-screen and cross-modality support, rather than independent patient-cohort validation. Finally, although the major functional associations were not attributable to a single major cancer lineage and no statistically supported modification by recorded donor sex was detected, these findings do not establish equivalence among lineages or sexes. In particular, recorded donor sex in established cell lines is not a surrogate for current endocrine exposure or androgen receptor signaling. In breast cancer prognosis studies, proliferation-related expression has also contributed to apparent prognostic associations of biologically unrelated signatures^64^, illustrating the need to distinguish state-specific information from shared transcriptional confounders.

Future studies can address these limitations at several levels. Direct experimental comparison of models with high and low ASTS scores will be required to determine which mitochondrial functions underlie the observed dependency and whether signaling dependence and mitochondrial vulnerability are mechanistically related. Longitudinal models of EGFR or MAPK pathway inhibition could determine whether changes in ASTS precede, accompany, or occur independently of the acquisition of drug resistance. Evaluation in independent patient cohorts with tumor transcriptomic and treatment-response data will also be necessary before ASTS can be considered for clinical stratification. Such studies may ultimately determine whether ASTS itself has predictive value or instead serves primarily as a route to identifying the molecular programs responsible for the associated vulnerabilities.

Taken together, the broader significance of ASTS lies less in any single drug association than in the strategy by which the functional phenotypes were uncovered. A transcriptional state derived from a defined in vivo hormonal perturbation and fixed independently of the downstream functional datasets remained informative after projection into an unrelated human cancer-cell context. In that setting, ASTS identified associations with signaling dependence and mitochondrial vulnerability that were not used in its construction and were not fully explained by the canonical oncogenic driver variables examined here. The present study therefore provides a proof of concept that transcriptional state information can complement genomic characterization when functional-genomic resources are used to prioritize candidate therapeutic vulnerabilities. Whether this principle extends to other endocrine or physiological perturbations, and ultimately to patient-level treatment stratification, will require systematic validation.

## Data availability

Raw mouse adrenal zona fasciculata RNA-seq data are available from the DNA Data Bank of Japan Sequence Read Archive under accession numbers DRA015952 and DRA017476. Processed mouse RNA-seq data, frozen ASTS signatures, ASTS scores, and derived pharmacogenomic, CRISPR dependency, and robustness results supporting this study are available in Zenodo (version 1.0.0) at https://doi.org/10.5281/zenodo.2317814965. Third-party source datasets from DepMap, PRISM, GDSC2, CTRPv2, and MSigDB are not redistributed and should be obtained from their original providers.

## Code availability

Analysis code and supporting reproducibility materials are available at GitHub (takbaba-lab/ASTS-cancer-dependency). The manuscript-associated software release, version V1.0.1, is archived in Zenodo at https://doi.org/10.5281/zenodo.2335504866.

## Acknowledgement

This work was supported by JSPS KAKENHI Grant Numbers 23K07991, 25K24532, and 26H02340. The computation was carried out using the computer resource offered under the category of General Projects by Research Institute for Information Technology, Kyushu University.

## Author Contributions

Y.M. and T.B. conceived and designed the study, performed the analyses, interpreted the results, and wrote the manuscript. Both authors reviewed and approved the final manuscript.

## Supplementary Information

### Supplementary figures and legends

Figure S1. Comparison of standard and ERCC-based normalization of mouse adrenal zF RNA-seq data.

*Figure: page 2; legend: page 3*

Figure S2. Construction and mouse-to-human projection of the ASTS signature.

*Figure: page 4; legend: page 5*

Figure S3. Robustness of transcriptomic associations with ASTS to cell-cycle and canonical driver adjustment.

*Figure: page 6; legend: page 7*

Figure S4. Robustness and pathway specificity of ASTS-associated drug-response phenotypes.

*Figure: page 8; legend: page 9*

Figure S5. Specificity and pharmacological support of the mitochondrial dependency associated with ASTS.

*Figure: page 10; legend: page 11*

Figure S6. Lineage robustness and donor-sex analyses of ASTS-associated functional phenotypes.

*Figure: page 12; legend: page 13*

---

**Figure S1.**
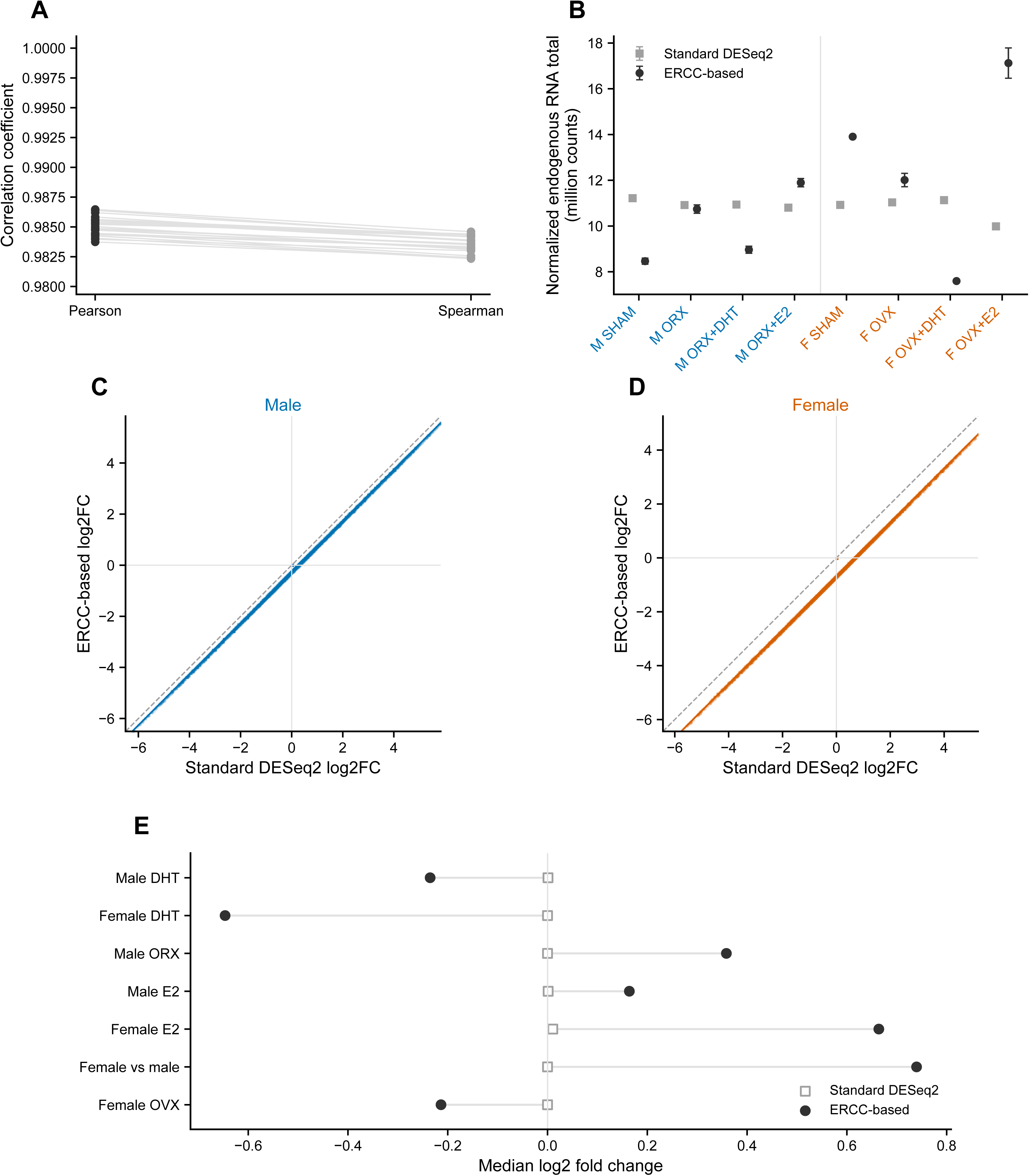
Comparison of standard and ERCC-based normalization of mouse adrenal zF RNA-seq data. **(A)** Agreement between the two counting procedures used for the RNA-seq dataset. Pearson and Spearman correlation coefficients were calculated for each sample, with paired values connected by lines. Mean Pearson and Spearman correlation coefficients were 0.985 and 0.984, respectively. Lines pair the two coefficients for each of the 24 samples; they are not confidence intervals. **(B)** Group-wise endogenous RNA abundance estimated using standard DESeq2 normalization and ERCC-based normalization. Points show group means and error bars indicate standard deviations (SD); n = 3 biological replicates per group. Male group labels are blue and female group labels are orange; gray squares and black circles indicate standard DESeq2 and ERCC-based normalization, respectively. M SHAM, sham-operated male; M ORX, orchiectomized male; M ORX+DHT, DHT-treated orchiectomized male; M ORX+E2, E2-treated orchiectomized male; F SHAM, sham-operated female; F OVX, ovariectomized female; F OVX+DHT, DHT-treated ovariectomized female; F OVX+E2, E2-treated ovariectomized female. Values are expressed in millions of normalized counts. **(C, D)** Gene-wise comparison of DHT-induced log2 fold changes obtained by standard DESeq2 and ERCC-based normalization in male **(C)** and female **(D)** zF cells. The dashed line indicates identity; colored solid lines indicate identity shifted by the median ERCC-minus-standard difference. Horizontal and vertical reference lines indicate zero. **(E)** Median log2 fold changes for the indicated hormonal and sex contrasts obtained using standard DESeq2 normalization (open squares) and ERCC-based normalization (filled circles). ERCC-based normalization retained the global component of the RNA-output response, whereas standard DESeq2 normalization centered this component and primarily captured relative transcriptional remodeling.

**Figure S2.**
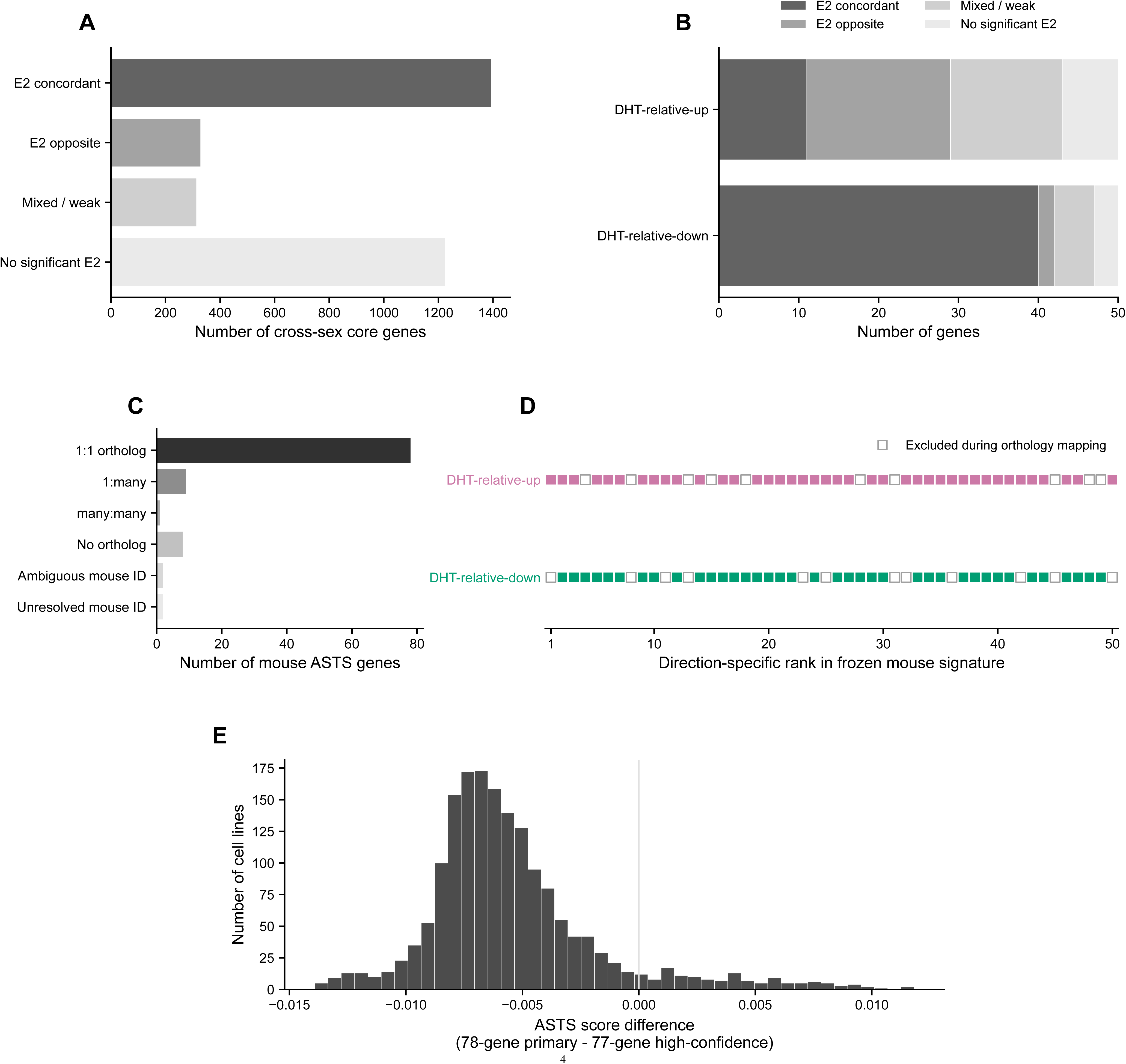
Construction and mouse-to-human projection of the ASTS signature. **(A)** Classification of the 3,259 cross-sex DHT-responsive core genes according to their response to E2. Genes were classified as E2 concordant, E2 opposite, mixed/weak, or without a significant E2 response. **(B)** E2-response composition of the frozen 100-gene mouse ASTS, shown separately for the 50 DHT-relative-up and 50 DHT-relative-down genes. The signature was frozen before pharmacogenomic analyses. **(C)** Mouse-to-human orthology mapping of the 100 frozen mouse ASTS genes. Seventy-eight genes mapped to one-to-one human orthologs and were retained in the primary human ASTS. **(D)** Retention of individual DHT-relative-up and DHT-relative-down genes after mouse-to-human orthology mapping, ordered by their direction-specific rank in the frozen mouse signature. Filled symbols indicate retained one-to-one orthologs, whereas open symbols indicate genes excluded during orthology mapping. **(E)** Sensitivity of DepMap ASTS scores to exclusion of the single lower-confidence one-to-one ortholog. The histogram shows the difference between scores calculated with the 78-gene primary signature and the 77-gene high-confidence sensitivity set across 1,719 cell lines. The two scores were highly concordant (Spearman ρ = 0.995); the median and maximum absolute score differences were 0.0063 and 0.0139, respectively. The solid vertical line marks zero.

**Figure S3.**
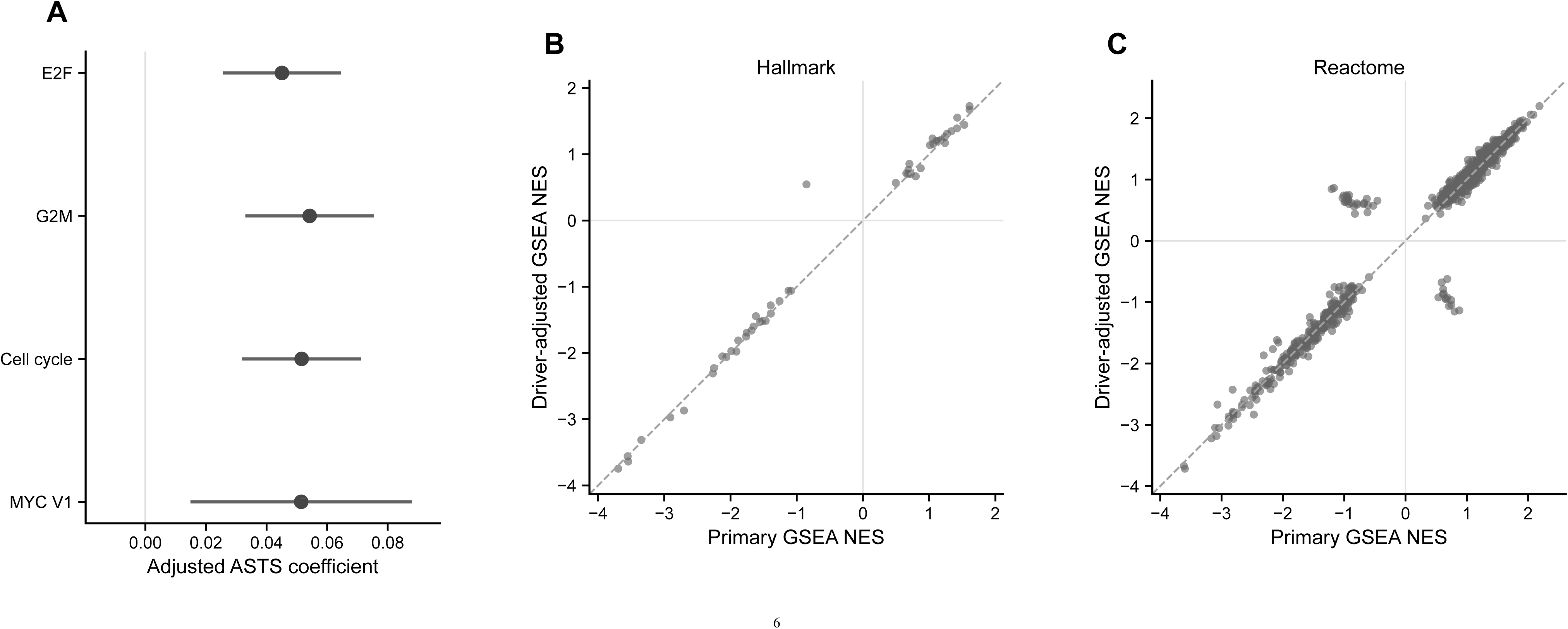
Robustness of transcriptomic associations with ASTS to cell-cycle and canonical driver adjustment. **(A)** Associations between ASTS and independently derived E2F, G2M, combined cell-cycle, and MYC V1 transcriptional scores. Points indicate standardized ASTS coefficients and horizontal lines indicate 95% confidence intervals from multivariable models including lineage, donor sex, proliferation, and canonical oncogenic driver covariates. Genes belonging to the ASTS signature were excluded when constructing these pathway scores. Each model included 1,636 cell lines. ASTS and the outcome score were z-standardized within the analyzed samples; confidence intervals were calculated as β ± 1.96 × HC3 SE. **(B)** Comparison of Hallmark GSEA normalized enrichment scores (NESs) obtained from the primary transcriptome-wide ASTS association analysis and from the corresponding driver-adjusted analysis. The two analyses showed strong concordance (Spearman ρ = 0.994; direction concordance = 98.0%). **(C)** Corresponding comparison for Reactome pathways (Spearman ρ = 0.985; direction concordance = 96.6%). These analyses indicate that the major transcriptomic architecture associated with ASTS was largely preserved after adjustment for canonical driver alterations. In panels B and C, dashed diagonals indicate identity and horizontal and vertical reference lines indicate zero.

**Figure S4.**
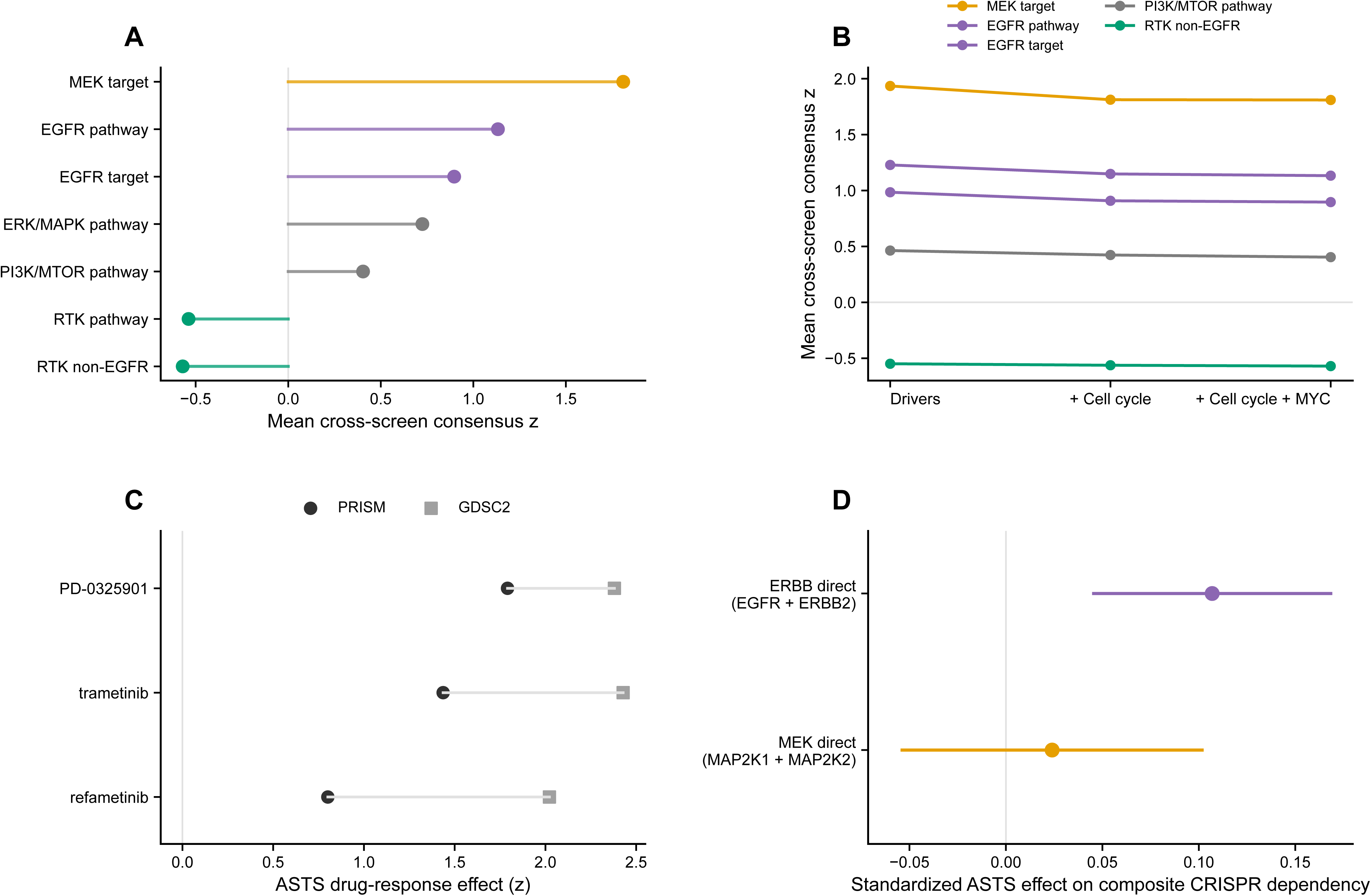
Robustness and pathway specificity of ASTS-associated drug-response phenotypes. **(A)** Mean cross-screen consensus z scores for predefined drug classes after adjustment for canonical driver alterations, cell-cycle activity, and MYC activity. Positive centered consensus z scores indicate effects shifted toward resistance relative to the screen-specific means of the matched drugs. The signs of the original regression coefficients determine whether ASTS is associated with reduced or increased drug sensitivity. **(B)** Stability of mean cross-screen consensus z scores across sequential adjustment models including canonical drivers alone, drivers plus cell-cycle activity, and drivers plus cell-cycle and MYC activity. The class-level shifts toward higher centered consensus z scores for MEK- and EGFR-directed drugs and lower scores for non-EGFR RTK inhibitors were retained after additional adjustment. **(C)** Screen-specific ASTS effects for the MEK inhibitors PD-0325901, trametinib, and refametinib in PRISM and GDSC2. Coefficients were centered and scaled across matched drugs within each screen; positive z scores indicate effects above the corresponding screen-specific mean. **(D)** Composite CRISPR dependency associations for the ERBB-direct axis (EGFR and ERBB2) and the MEK-direct axis (MAP2K1 and MAP2K2). Points indicate standardized ASTS coefficients and horizontal lines indicate 95% confidence intervals from the stringent multivariable model. Positive values indicate reduced dependency in ASTS-high cell lines. The ERBB-direct dependency association was evident, whereas the MEK-direct composite showed little evidence of reduced dependency, distinguishing the pharmacological MEK-resistance phenotype from direct MAP2K1/2 dependency loss. FDR values were corrected across three composite axes; the ERBB-direct axis had P = 0.000650 and FDR = 0.00195.

**Figure S5.**
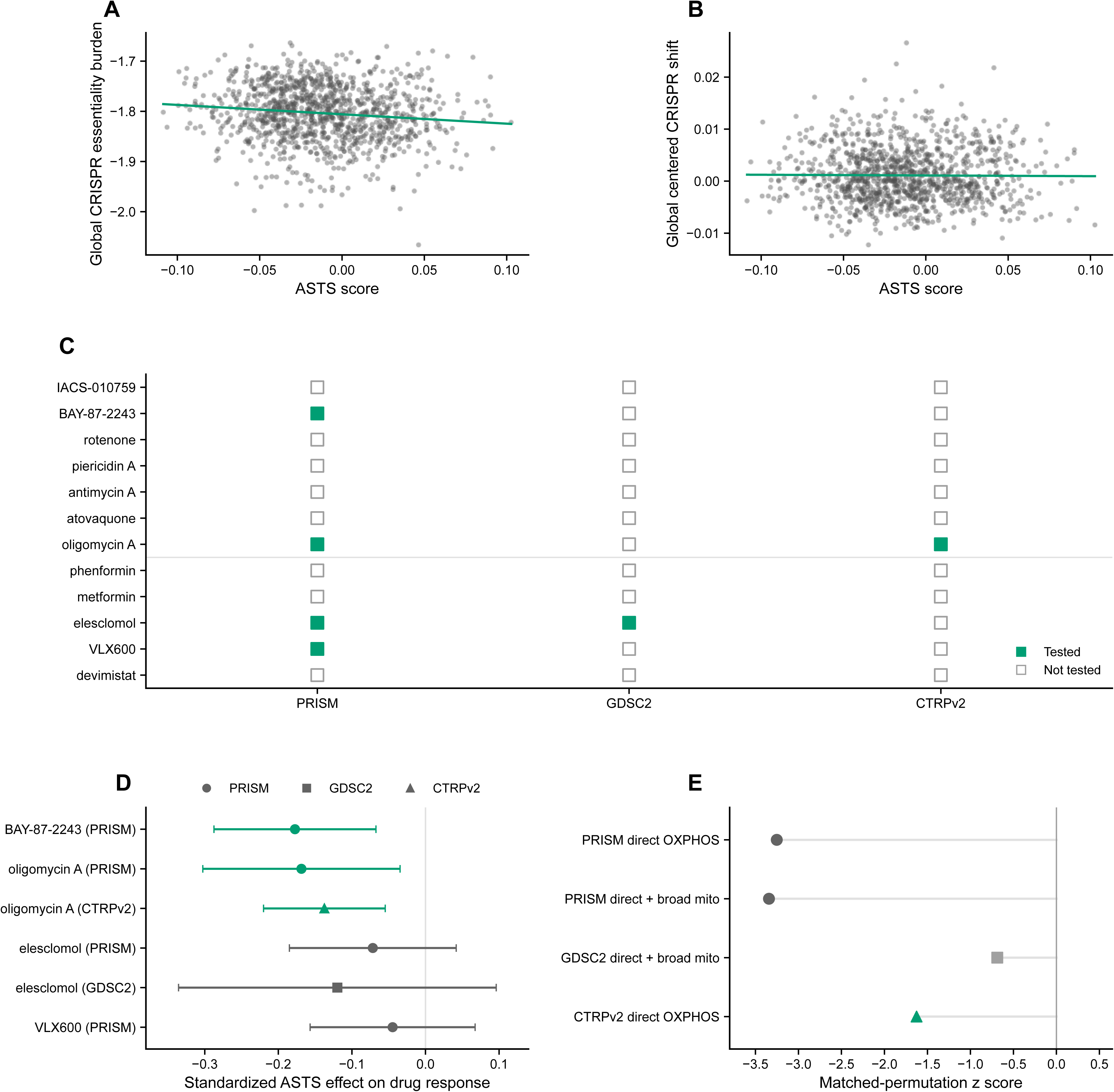
Specificity and pharmacological support of the mitochondrial dependency associated with ASTS. **(A)** Relationship between ASTS score and a global CRISPR essentiality-burden metric calculated from non-mitochondrial common-essential genes. A weak negative association was observed (n = 1,135; Spearman ρ = −0.134, P = 6.28 × 10^-6). **(B)** Relationship between ASTS score and the global gene-centered CRISPR shift. No appreciable association was observed (n = 1,135; Spearman ρ = −0.011, P = 0.708). Together with explicit burden adjustment in the genome-wide dependency models, these analyses distinguish the mitochondrial phenotype from a generalized shift in CRISPR essentiality. **(C)** Availability of the mechanism-defined mitochondrial drug candidates across PRISM, GDSC2, and CTRPv2. Green filled squares indicate available compounds and gray open squares indicate compounds not found. Evaluable sample sizes are provided in Source Data. Candidates were grouped into Tier 1 direct OXPHOS perturbations and Tier 2 broader mitochondrial perturbations. OXPHOS, oxidative phosphorylation. **(D)** Screen-specific standardized ASTS effects for all mechanism-defined mitochondrial candidates available for analysis. Points indicate standardized regression coefficients and horizontal lines indicate 95% confidence intervals. Circles, squares, and triangles denote PRISM, GDSC2, and CTRPv2, respectively. Negative values indicate greater drug sensitivity in ASTS-high cell lines. Direct OXPHOS perturbations are green; broader mitochondrial perturbations are gray. Oligomycin A showed concordant negative effects in PRISM and CTRPv2. **(E)** Matched-drug permutation analysis. Negative z scores indicate candidate-set mean coefficients below the matched null expectation; their signs alone do not establish increased sensitivity (original coefficients, panel D). PRISM direct OXPHOS and direct-plus-broader sets had empirical one-sided P = 0.00210 and 0.00140, respectively (both FDR = 0.00315). These two tests and the GDSC2 single-candidate test were corrected jointly; GDSC2 had P/FDR = 0.239. The two CTRPv2 sets both contained only oligomycin A and had P/FDR = 0.0200 after separate correction. Limited coverage supports pharmacological follow-up rather than broad OXPHOS-inhibitor sensitivity. Circles, squares, and triangles denote PRISM, GDSC2, and CTRPv2, respectively.

**Figure S6.**
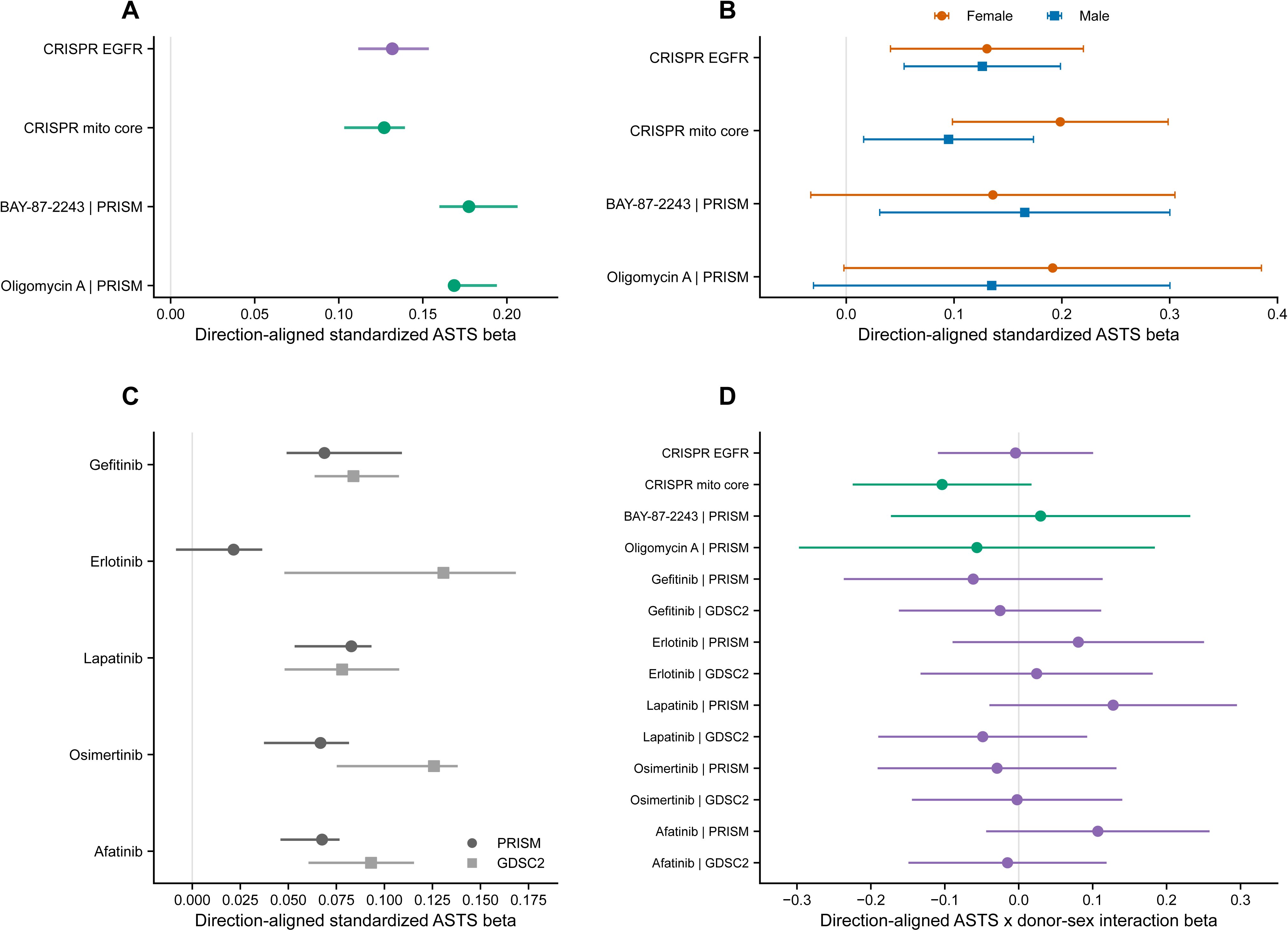
Lineage robustness and donor-sex analyses of ASTS-associated functional phenotypes. (A) Leave-one-lineage-out (LOLO) analysis of the four primary endpoints. Points show direction-aligned standardized full-model ASTS effects; horizontal lines show the minimum-to-maximum range after lineage exclusion, rather than confidence intervals. Positive values support the predefined phenotype. EGFR and mitochondrial-core dependency were evaluated across 18 exclusions each; BAY-87-2243 and oligomycin A across five and three, respectively. Purple denotes EGFR; green denotes mitochondrial endpoints. (B) Female-specific (orange circles) and male-specific (blue squares) ASTS slopes from the combined ASTS-by-donor-sex interaction model. Horizontal lines indicate 95% confidence intervals. Both CRISPR endpoints included 1,082 cell lines (465 female, 617 male); BAY-87-2243 included 438 (190 female, 248 male); oligomycin A included 342 (151 female, 191 male). Models included lineage, proliferation, canonical drivers, cell-cycle and MYC V1 scores. Mitochondrial-core models omitted global CRISPR burden covariates. (C) LOLO analysis of five EGFR inhibitors in PRISM (circles) and GDSC2 (squares). Symbols show full-model standardized effects; lines show the minimum-to-maximum range after lineage exclusion. Positive values indicate ASTS-high resistance. (D) Direction-aligned ASTS-by-donor-sex interaction coefficients are shown for the four primary functional endpoints and EGFR-inhibitor responses in PRISM and GDSC2. Points indicate interaction coefficients and horizontal lines indicate 95% confidence intervals. Negative values indicate a stronger predefined phenotype in female-derived cell lines, whereas positive values indicate a stronger phenotype in male-derived cell lines. No interaction remained significant after joint Benjamini-Hochberg correction across the 14 endpoints (minimum FDR = 0.751). Thus, no statistically supported donor-sex effect modification was detected, although this analysis does not establish equivalence between sexes. Donor sex denotes the recorded sex of the individual from whom the cell line was derived and does not represent androgen exposure. Purple denotes EGFR-related and green mitochondrial endpoints. Vertical reference lines in all panels indicate zero.

